# Extracellular vesicles and high-density lipoproteins: Exercise and estrogen-responsive small RNA carriers

**DOI:** 10.1101/2022.02.28.482100

**Authors:** Sira Karvinen, Tia-Marje Korhonen, Tero Sievänen, Jari E. Karppinen, Hanna-Kaarina Juppi, Veera Jakoaho, Urho M. Kujala, Jari A. Laukkanen, Maarit Lehti, Eija K. Laakkonen

## Abstract

Decreased systemic estrogen levels (i.e., menopause) affect metabolic health. However, the detailed mechanisms underlying this process remain unclear. Both estrogens and exercise have been shown to improve metabolic health, which may be partly mediated by circulating microRNA (c-miR) signaling. In recent years, extracellular vesicles (EV) have increased interest in the field of tissue crosstalk. However, in many studies on EV-carried miRs, the co-isolation of high-density lipoprotein (HDL) particles with EVs has not been considered, potentially affecting the results. Here, we demonstrate that EV and HDL particles have distinct small RNA (sRNA) content, including both host and nonhost sRNAs. Exercise caused an acute increase in relative miR abundancy in EVs, whereas in HDL particles, it caused an increase in transfer RNA-derived sRNA. Furthermore, we demonstrate that estrogen deficiency caused by menopause blunts acute exercise-induced systemic miR-response in both EV and HDL particles.

**HIGHLIGHTS:**

- Extracellular vesicles and HDL particles have a distinct sRNA content
- Extracellular vesicles and HDL particles carry both host and nonhost sRNA cargo
- Estrogen deficiency blunts the c-miR-response induced by acute exercise
- Exercise responsive miRs in HT users may regulate the choice of energy substrate

## INTRODUCTION

Women experience an increment in metabolic and cardiovascular disease risks after the onset of menopause (Clegg et al., 2017). Especially the decrease in systemic estrogen levels have been associated with unfavorable changes in metabolic health (Carr, 2003). Both estrogens and exercise have been shown to improve cardiometabolic health by enhancing mitochondrial function and decreasing inflammation (Torres et al., 2018; Vieira-Potter et al., 2015). However, we have shown that physical activity does not entirely offset the unfavorable changes in lipid profile associated with the menopausal transition (Hyvärinen et al., 2021; Karvinen et al., 2019). Discovering the route and mechanisms via which estrogens alter metabolism is crucial to understanding the physiological characteristics of menopause that compromise women’s cardiometabolic health.

Recently, small non-coding RNAs (sRNAs), including microRNAs (miRs), have been recognized as a part of the regulatory network governing gene expression (Gomes et al., 2018). miRs are gene regulators that typically repress protein synthesis, affecting numerous biological processes, such as body metabolism, inflammation, apoptosis, and aging (Subramanian and Steer, 2010; Zhang et al., 2015a). In the context of exercise, circulating miRs (c-miRs) have been shown to mediate exercise adaptations (Sapp et al., 2017). In fact, c-miR expression rapidly changes the response to a changing environment, which makes c-miRs attractive biomarkers to use in studying exercise responses (Bartel, 2009; Friedman et al., 2009).

C-miRs are transported in the blood circulation via several types of vesicular or protein-based particles (Sapp et al., 2017). Of these carrier particles, the role of extracellular vesicles (EVs) has attracted particular interest in the field of tissue crosstalk during exercise (Whitham et al., 2018) because an acute bout of exercise transiently changes the miR content of EVs (Sapp et al., 2017). We have previously demonstrated that the circulating concentration of estrogen is another factor associated with differences in the c-miR content of EVs in women (Kangas et al., 2017). We have also shown that estrogen-responsive c-miRs are linked to several clinical biomarkers of metabolic health. To our knowledge, the question of whether estrogen or menopausal status affects exercise-induced changes in c-miR expression, potentially affecting women’s metabolic responses to acute exercise, has not been studied previously.

In most of earlier studies of EV-carried miRs, the question of whether high-density lipoprotein (HDL) particles, which are very abundant in blood plasma, may be co-isolated with EVs has not been considered (Simonsen, 2017). This occurs, for example, when EVs are isolated using methods that are simply based on the size or density of the particle. The potential co-isolation of HDL with EVs is important to consider because, in addition to acting in reverse cholesterol transport, HDL particles have been proven to carry miRs and other sRNA species, which may also participate in tissue crosstalk (Vickers and Michell, 2021). HDL particles are especially interesting in the context of menopause because, after menopausal transition, both HDL cholesterol (HDL-C) level and HDL particle number increase, but cardiovascular protection deteriorates (Khoudary et al., 2016, 2018). Hence, it is plausible to hypothesize that HDL function and/or composition are affected by menopause-associated estrogen deficiency. However, the sRNA content of these two major extracellular RNA-carriers, EV and HDL particles, has not been fully characterized or compared before.

Here, we demonstrate that EV and HDL particles have distinct sRNA content. In addition to sRNA of human (host) origin, we found both to carry a substantial amount of nonhost sRNA. EVs had a relatively larger amount of host sRNA as compared with HDL particles. Of the host sRNA, miRs were the most numerous RNA species present in EVs, whereas in HDL particles, the most abundant species were ribosomal RNA (rRNA) -derived sRNAs (rDRs). We further investigated whether an acute bout of exercise induces changes in sRNA content carried via EV or HDL particles. We observed an increase in the relative miR abundancy in EVs, whereas in HDL particles, we noticed an increase in transfer RNA (tRNA) -derived sRNAs (tDRs), while miRs and rDRs became less abundant. By focusing on the miRs whose functions are best characterized, we also examined whether exercise response differs based on estrogen status. Strikingly, among estrogen-deficient postmenopausal women, no exercise response was detected. Our results highlight that the estrogen deficiency caused by menopause diminishes the acute exercise-induced miR-response in both EV and HDL particles.

## RESULTS

### EV and HDL particles have distinct sRNA content

To investigate the RNA content of EV and HDL particles, we collected plasma samples after overnight fasting from a subset of “Estrogen, microRNAs and the risk of metabolic dysfunction (EsmiRs) study” (Laakkonen et al., 2021) participants, consisting of 18 postmenopausal women (age range 52–57 years, Table S1). Of these women, nine used estrogen-based hormonal therapy (HT), and nine were HT nonusers.

We isolated plasma EV and HDL particles via sequential ultracentrifugation, followed by size-exclusion chromatography (SEC), allowing the separation of these sRNA-carrying extracellular particles (Simonsen, 2017). After validating successful isolation, resulting in a clear separation of EV and HDL particles with Western blot (WB), dot blot (DB), electron microscopy (EM), and immunoelectron microscopy (IEM) analysis (Figures 1A-F), we proceeded with RNA isolation and Next-Generation Sequencing (NGS) analysis with purified EV and HDL samples obtained from 14 postmenopausal women including seven HT users (Table S1).

**Figure 1.**
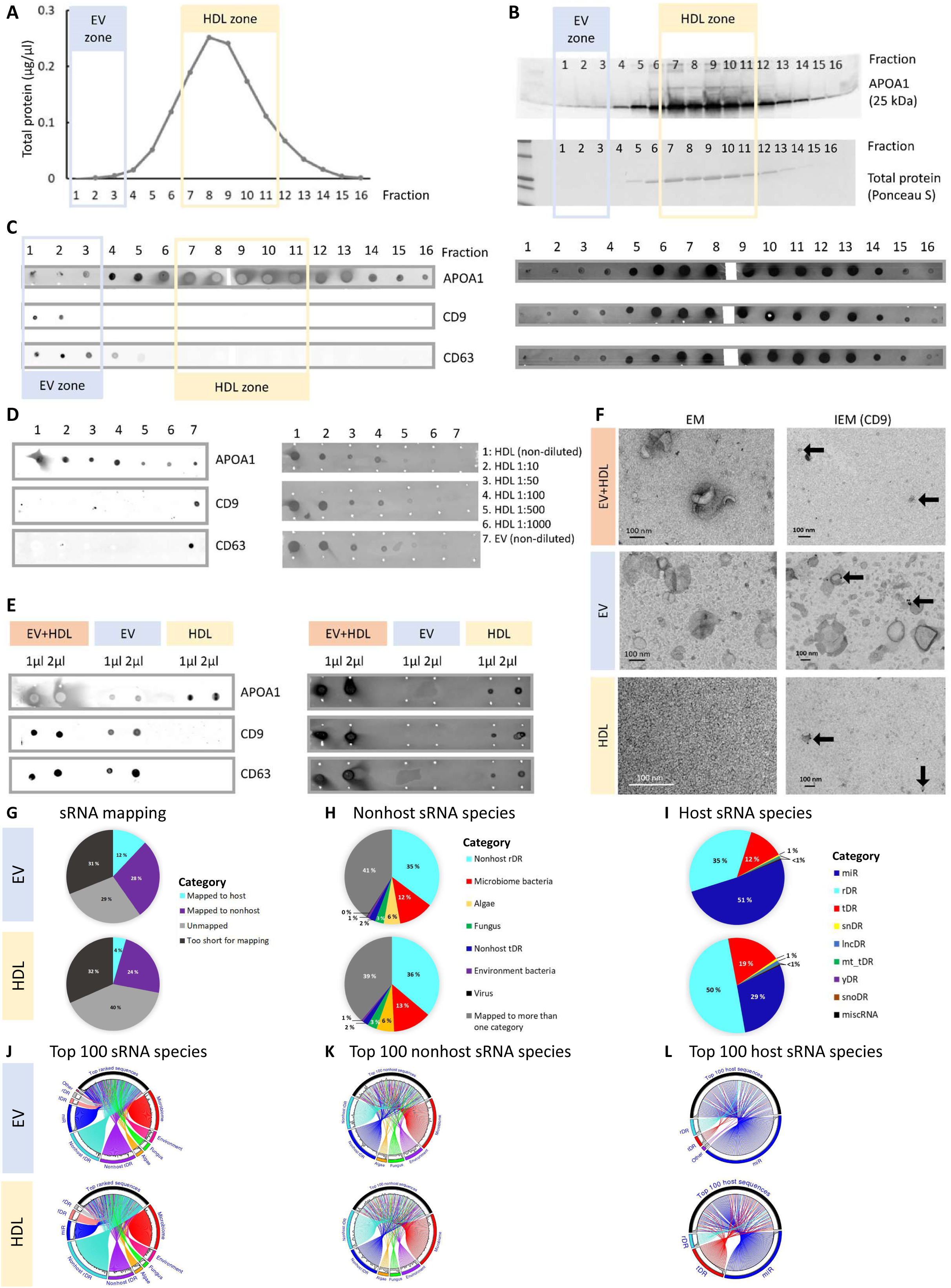
EV and HDL particles have distinct sRNA content. **(A)** Total protein (BCA) analysis of SEC fractions 1–16. **(B)** Verification of HDL particle enrichment in fractions 7–11 via WB using APOA1 antibody and corresponding total protein blot stained with PonceauS. **(C)** DB verification of enrichment of HDL particles into fractions 7–11 with APOA1 antibody and enrichment of EVs into fractions 1–3 with CD9 and CD63 antibodies (left panel) and corresponding total protein blots stained with PonceauS (right panel). **(D)** HDL dilution series and non-diluted EV sample as positive control when visualizing APOA1, CD9, and CD63 antibodies (left panel) and corresponding total protein blots stained with PonceauS (right panel). **(E)** Fractions EV+HDL, and EV and HDL separated via SEC and visualized with 1µl and 2µl sample with APOA1, CD9, and CD63 antibodies (left panel) and corresponding total protein blots stained with PonceauS (right panel). **(F)** Representative EM and IEM (stained with CD9) images of fractions EV+HDL, EV, and HDL. **(G)** sRNA categories in EV and HDL particles. **(H)** Non-host sRNA categories in EV and HDL particles. **(I)** Host sRNA categories in EV and HDL particles. **(J)** Top 100 sRNA species in EV and HDL particles. **(K)** Top 100 nonhost sRNA species in EV and HDL particles. **(L)** Top 100 host sRNA species in EV and HDL particles.

The analyzed EV samples had an average of 4.18 million raw reads per sample, and the HDL samples had an average of 4.50 million raw reads/sample. The maximum number of reads in a sample was over 11 million, and the minimum was 659,000. Two HDL samples that had less than 600,000 reads were excluded from the analysis. We utilized the “Tools for Integrative Genome analysis of Extracellular sRNAs (TIGER)” data analysis pipeline (Allen et al., 2018), allowing for the comprehensive investigation of the acquired sRNA sequences. After adapter trimming, there was on average 919,000 sRNA reads in EV and 920,000 sRNA reads in HDL pre-exercise samples. Similar to Allen *et al*., relatively large amount of all sequences remained unmapped or were deemed too short for reliable mapping (<16 nucleotides). Out of all sRNA sequences, 12.0% of EV-carried and 4.5% of HDL-carried were shown to have a human origin (Figure 1G, Table S2), and 28.2% and 23.5% respectively had a nonhost origin. Supporting our findings, Wang *et al*. (2012) observed that human plasma contains a significant amount (∼40%) of exogenous RNA, whereas Allen *et al*. (2018) showed, in isolated mouse HDL, that only a relatively small amount of sRNA was mapped to host-origin (Allen et al., 2018; Wang et al., 2012). To our knowledge, a similar examination of isolated EVs has not been carried out previously.

The relative abundances of sRNAs of nonhost origin followed similar trends in EV and HDL particles. Most of the sRNAs were mapped to more than one category (41.0% in EVs and 38.8% in HDL particles). The second largest group was nonhost rDRs (35.2% and 36.1%, respectively) (Figure 1H, Table S2). Interestingly, 12.0% of EV-carried and 13.1% of HDL-carried sRNA was mapped to the bacterial microbiome. Supporting our findings, Wang *et al*. (2012) suggested the possibility that exogenous RNA found in plasma may mediate human-microbiome interaction (Wang et al., 2012). The sRNA of algal, fungal, environmental bacterial, and viral origin formed the smallest relative fractions in both EV (≤5.6%) and HDL particles (≤6.0%) (Figure 1H, Table S2).

Of the host sRNA, miRs were the most numerous RNA species present in EVs (51.2%), highlighting the role of EVs as mediators of miR-signaling. The next most common sRNA species in EVs were rDRs (34.8%) and tDRss (12.1%) (Figure 1I, Table S2). Similarly, Huang *et al*. (2013) observed that miRs and rRNAs were the most abundant RNA species in EVs isolated from human plasma (Huang et al., 2013). In contrast to EVs, in HDL particles, rDRs were the most numerous sRNA species (49.9%), followed by miRs (29.3%) and tDRs (18.5%) (Figure 1I, Table S2). Michell *et al*. (2016) also found that isolated human HDL carried different sRNA species, but in their analysis, the largest classes were miRs and tDRs (38% and 37%, respectively), followed by miscellaneous RNAs (miscRNAs) and rDRs (11% and 10%, respectively) (Michell et al., 2016). In our samples, low-abundance sRNA species, including long non-coding (lncRNA) -derived sRNAs (lncDRs), small nuclear RNA (snRNA) -derived sRNAs (snDRs), small nucleolar RNA (snoRNA) -derived sRNAs (snoDRs), mitochondrial tDRs, yRNAs (yDRs), and miscRNA -derived sRNAs (miscDRs), formed 1.5% of EV-carried and 2.3% of HDL-carried sRNA (Figure 1I, Table S2). Previous studies have demonstrated that sRNA species that originate from tRNAs, rRNAs, and snoRNAs may possess miR-like activity, altering gene expression (Ender et al., 2008; Lee et al., 2009; Li et al., 2012; Wei et al., 2013).

Of the top hundred sRNA species present in EVs, most belonged to the microbiome, nonhost rDR, tDR, and miR classes (Figure 1J). In HDL, the top hundred sequences belonged to the nonhost rDR and microbiome classes, followed by the nonhost tDR and miR classes (Figure 1J). The observation that the most abundant sequences of all sRNA species in both EV and HDL particles were mapped to the microbiome further highlight the potential role of nonhost RNA in mediating the human-microbiome interaction (Wang et al., 2012). Accordingly, of the top hundred sRNA sequences of nonhost origin, the majority belonged to the microbiome in both EV and HDL particles, followed by rDR and tDR (Figure 1K). The top hundred host sRNA species were miRs in both EV and HDL particles, followed by rDRss and tDR in EVs, whereas in HDL particles, tDRs were more common than rDRs (Figure 1L). These results further confirm that EV and HDL particles have distinct sRNA content, with EVs carrying mainly miRs, whereas HDL particles also carry a substantial number of host tDRs, in addition to rDRs and miRs.

### An acute bout of exercise induces a distinct change in the sRNA content of EV and HDL particles

To examine the change in the sRNA content of EVs and HDL in response to acute exercise, we harvested blood plasma samples after overnight fasting prior to (PRE), immediately after (POST) and 1 hour after (1h POST) a bicycle ergometer VO_2peak_ test, and TIGER analysis was performed for all samples.

In the EVs, we observed a small increase in the relative abundance of miRs (+3.1% points) and tDRs (+2.1% points), while the relative amount of rDRs decreased (−5.0% points) after exercise (POST-PRE) (Figure 2A, Table S2). In response to exercise stimulus, our results showed a transient increase in miRs, because there was an -7.6% point decrease in the relative miR content 1h after exercise (1h POST-POST). In contrast, in HDL, we observed a small decrease in the relative abundance of miRs (−0.9% points), while the relative amount of tDRs increased (+13.7% points) and rDRs decreased (− 12.9% points) markedly immediately after exercise (POST-PRE). Also, in HDL, the relative amount of miRs continued to decrease 1h after exercise stimulus (−6.9% points), while the relative amount of rDRs increased (+11.1% points) (1h POST-POST) (Figure 2A, Table S2).

**Figure 2.**
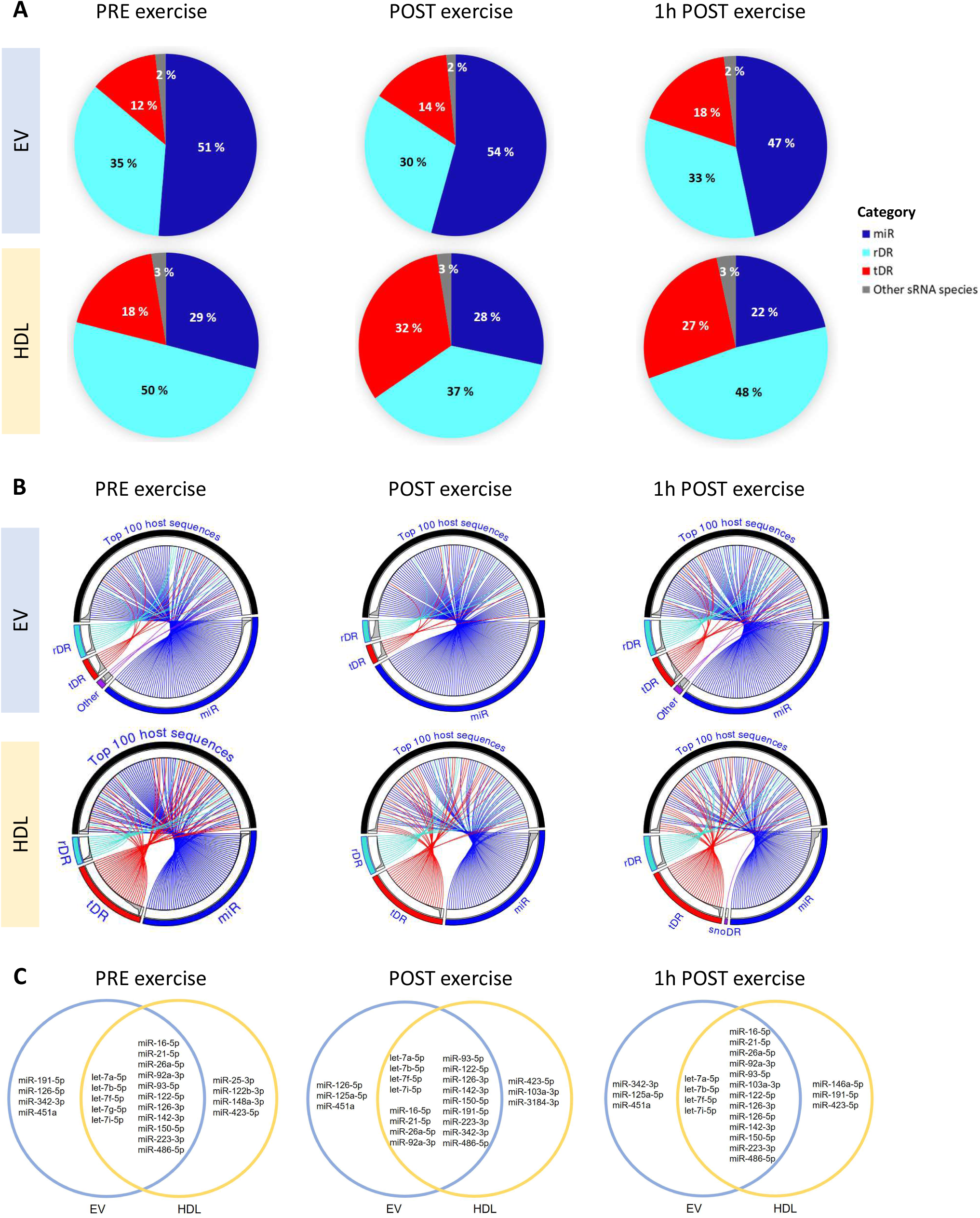
An acute bout of exercise induces a distinct change in the sRNA content of EV and HDL particles. **(A)** Relative % of sRNA species in carried in EV and HDL particles prior to exercise (PRE), immediately after exercise (POST), and 1 h after exercise (1h POST). **(B)** Top hundred host sRNA species in EV and HDL particles PRE, POST and 1h POST exercise. **(C)** Twenty most abundant miRs (NGS counts) present in EVs and HDL particles PRE, POST, and 1h POST exercise.

Considering the top hundred sequences in EVs, we observed that a larger number of sequences belonged to miRs immediately after exercise (POST) as compared with the PRE and 1h POST time points (Figure 2B). This observation further highlights that not only does the relative abundance of miRs increase after exercise, but miRs are also among the most abundant sRNA sequences found in EVs. The next most abundant sequences carried via EVs were rDRs and tDRs. Similarly to EVs, in HDL particles, the most abundant sRNA sequences were miRs at all three time points (Figure 2B). However, in HDL particles, tDRs were more commonly represented among the most abundant reads than in EVs (Figure 2B). Interestingly, in HDL, snoDRs were also observed among the top hundred sequences 1h POST exercise. Our results show that an acute bout of exercise induces a distinct change in the sRNA content of EV and HDL particles.

We examined DESeq2-normalized miRs (Figures S1A-B, Table S3) at each time point in EVs and HDL particles, and the twenty most abundant (highest number of counts) were included in the Venn diagrams (Figure 2C). The Venn diagrams show that, at each time point, the majority (16–17) of the most abundant miRs were common between EV and HDL particles (Figure 2C). Specifically, in EVs, miR-451a was among the twenty most abundant miRs at all three time points, whereas in HDL, a similar observation was made for miR-423-5p. Previously, the up-regulation of miR-451a has been shown in response to estrogen in mouse lymphocytes (Dai et al., 2008), while miR-423-5p is involved in ovarian function and estrogen secretion (Xie et al., 2020). Both miR-423-5p and 451a are up-regulated in the blood in response to exercise and may target the pathways coordinating metabolic regulation (Li et al., 2020; Olioso et al., 2019).

### Only HT users exhibited an acute exercise-induced c-miR response

Next, we investigated the potential differences in exercise response based on participant estrogen status. For that purpose, differential expression (DE) analysis was carried out to compare study groups composed of 14 EV and HDL samples from HT users and nonusers (n = 7/group). The study groups were matched for age, peak aerobic capacity (VO_2peak_), body mass index (BMI) and body fat percentage (Table S1). At this stage, we concentrated only on miRs. To visualize the miR response to acute exercise, we created miR trajectory figures showing up-regulated (red, FC ≥ 0.5) and down-regulated (blue, FC ≤ -0.5) miRs when comparing POST vs. PRE levels (Figures 3A-B). In addition, trajectories marked with black represent miRs that changed when comparing 1h POST vs. POST levels. Although numerous EV- and HDL-carried miRs responded to exercise in both HT users and nonusers (Figures 3A-B), only in HT users were differentially expressed miRs observed (Figures 3C-D).

**Figure 3.**
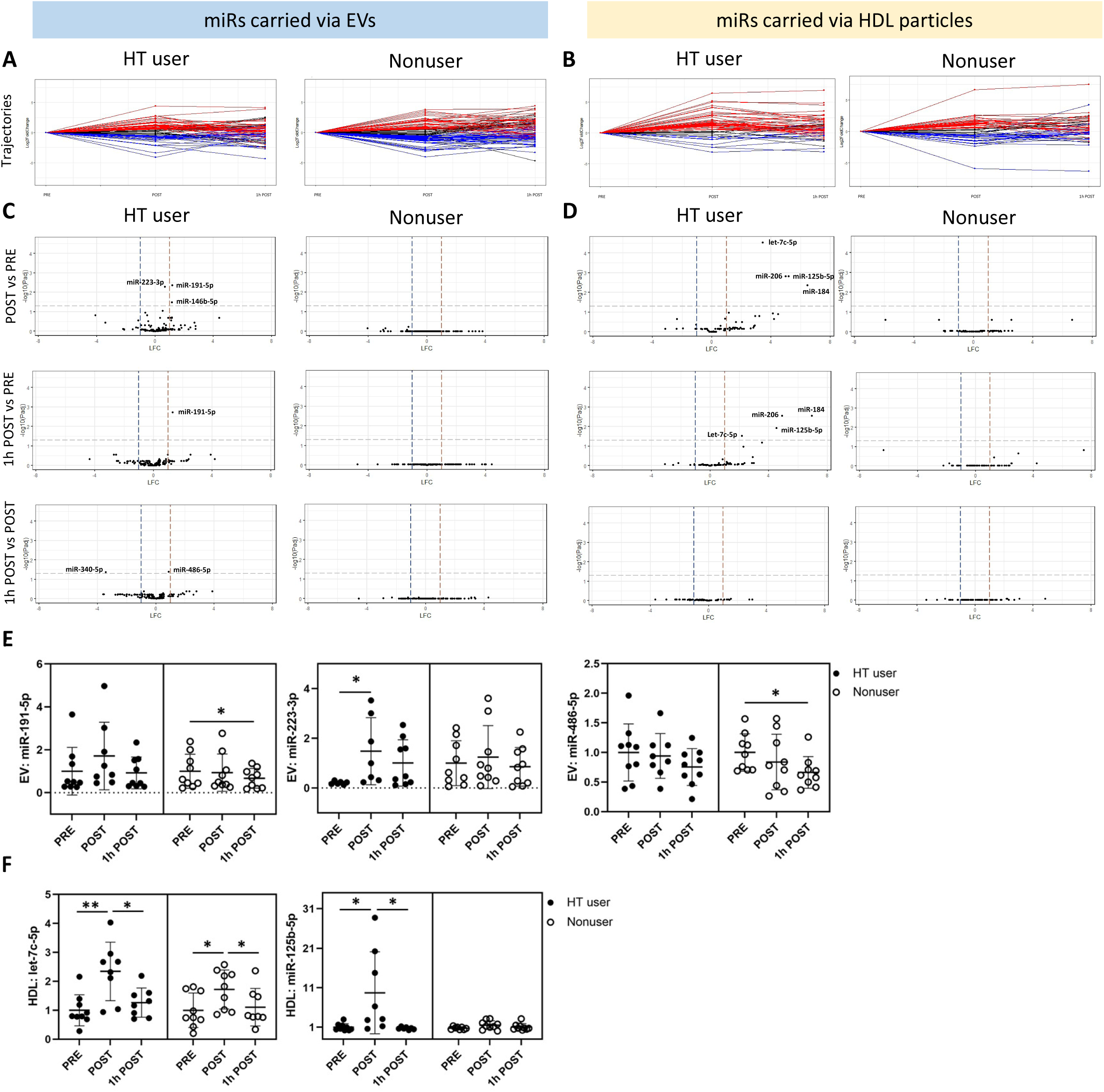
Only HT users exhibit an acute exercise-induced miR-response. **(A-B)** miR trajectories showing up-regulated miRs (RED) and down-regulated miRs (BLUE) in POST-to-PRE comparison and miRs whose expression changed (up- or down-regulation) in 1h POST-to-POST comparison (BLACK) **(C-D)** Volcano plots showing the differential expression of miRs between time points in EV and HDL particles in HT users and nonusers. **(E)** Significantly changed miRs in EVs in response to exercise, verified via qPCR. **(F)** Significantly changed miRs in HDL particles in response to exercise, verified via qPCR.

To identify differentially expressed miRs carried via EVs and HDL particles, we followed the statistical procedures of the DESeq2 R-package and used a paired samples *t*-test to compare exercise time points POST vs. PRE, 1h POST vs. PRE, and 1h POST vs. POST. The obtained results are visualized using volcano plots (Figures 3C–D), and numerical data are presented in Tables S4–7. In the POST vs. PRE comparison, miRs 191-5p, 223-3p, and 146b-5p were significantly up-regulated in the EVs of HT users (Figure 3C, Table S4). Of these, only miR-191-5p remained up-regulated at the time point 1h POST (Figure 3C, Table S4). In addition, when comparing 1h POST vs. POST, miR-486-5p was up-regulated, and miR-340-5p was down-regulated (Figure 3C, Table S4). Interestingly, there were no significant findings in the studied time points in nonusers (Figure 3C, Table S5).

In miRs isolated from HDL particles, we observed a significant up-regulation of miRs 125b-5p, 184, -206 and let-7c-5p in HT users when comparing time points POST vs. PRE, and all the miRs remained up-regulated at time point 1h POST (Figure 3D, Table S6). Again, we found no significant changes in the studied time points in nonusers (Figure 3D, Table S7).

We validated the significantly changed miRs with the highest expression in NGS with qPCR by using the complete cohort (n = 9/group) and confirmed the up-regulation of miR-223-3p in the EVs (Figure 3E) and let-7c-5p and miR-125b-5p in the HDL particles (Figure 3F) of the HT users. In contrast to the NGS results, we did observe some significant differences among nonusers via qPCR. In EVs, miR-191-5p and 486-5p were down-regulated when comparing 1h POST vs. PRE (Figure 3E), and in HDL, let-7c-5p was up-regulated in POST vs. PRE and down-regulated in 1h POST vs. POST comparisons (Figure 3F).

We carried out the same differential expression analysis with rDRs and tDRs (Figure S2C-F, Tables S8-15). In HT users, we did not observe differentially expressed rDR species in EVs POST vs. PRE comparison, but we did notice several changes in 1h POST vs. PRE and 1h POST vs. POST comparisons (Figure S2A, Table S8). The opposite was true for nonusers, who exhibited several differentially expressed rDR species in POST vs. PRE comparison, whereas no differences were observed in other comparisons (Figure S2A, Table S9). In HDL particles, we found no changes in rDRs in HT users, whereas in nonusers, there were several differentially expressed rDR species in HDL particles in POST vs. PRE and 1h POST vs. PRE comparisons (Figure S2B, Tables S10-11). Strikingly, all the rDR species that significantly changed in response to exercise in EVs and the majority of such species in HDL particles belonged to mitochondrial ribosomal subunits (Tables S8-11). Endurance exercise is known to increase mitochondrial biogenesis, as well as ribosome biogenesis in skeletal muscle (Mesquita et al., 2021), and ribosome biogenesis specifically has been associated with skeletal muscle hypertrophy (Figueiredo and McCarthy, 2019). Hence, we speculate that the observed changes in rDRs may reflect the response of muscle tissue to exhaustive exercise (VO_2peak_ test).

When examining tDRs in EVs, tRNA-Gln (anticodon TTG) was up-regulated in HT users in 1h POST vs. POST comparison (Figure S2C, Table S12). We observed no changes in nonusers (Figure S2C, Table S13). In HDL particles, two different tDRs coding for tRNA-Glu (anticodon TTC) were upregulated in HT users in 1h POST vs. PRE comparison (Figure S2D, Table S14). In nonusers, tRNA-Glu (anticodon TTC) was upregulated, and ten tDR species were downregulated in POST vs. PRE comparison. Additionally, tRNA-Glu (anticodon TTC) was upregulated when comparing 1h POST vs. PRE (Figure S2D, Table S15). The results highlight the fact that rDRs and tDRs have distinct responses to acute exercise in subjects differing in terms of their estrogen status, with rDRs showing more differentially expressed species.

### miRs responding to exercise in HT users may regulate the choice of energy substrate use

To understand the potential functional roles of the NGS-identified exercise-responsive c-miRs, we performed functional miR target analysis via miRNet (Chang et al., 2020) and miRPath v.3 (Vlachos et al., 2015). These analyses included all c-miRs (five from EVs and four from HDL particles) found to be differentially expressed in any of the time point comparisons (Figure 3C).

The miRNet analysis revealed that miRs changed in response to exercise in EVs are interconnected by sharing target proteins, which are regulated by two or more miRs (Figure 4A). Insulin-like growth factor 1 receptor (IGF1R) was the only target regulated by all five miRs; MAP kinase-interacting serine/threonine kinase 2 (MKNK2) was regulated by three miRs; and rho-related GTP-binding protein (RHOB), fork head box protein O1 (FOXO1), epithelial cell-transforming sequence 2 oncogene 2 (ECT2), and cystic fibrosis transmembrane conductance regulator (CFTR) were regulated by two miRs (Figure 4A). The analysis run via miRPath v.3 using the Kyoto Encyclopedia of Genes and Genomes (KEGG) pathways union option revealed ten significantly regulated pathways to be targeted by the five studied miRs, including the fatty acid degradation pathway, which is an important pathway for exercise responses (Figure 4B). Further analysis with miRPath v.3 using the genes union option revealed, in total, 28 KEGG signaling pathways that were significantly regulated by those five miRs and verified the interconnection of these miRs with FOXO signaling (Figure 4C, Table S16). FOXO1 is known to play a significant role in regulating whole-body energy metabolism. For example, it promotes the expression of gluconeogenic enzymes in the liver and has been suggested to promote the switch from using carbohydrates to using fatty acids as an energy source (Gross et al., 2009). Both insulin-like growth factor’s (IGF) and mitogen-activated protein kinase’s (MAPK) pathways interconnect through FOXO signaling pathways (Figure 4C).

**Figure 4.**
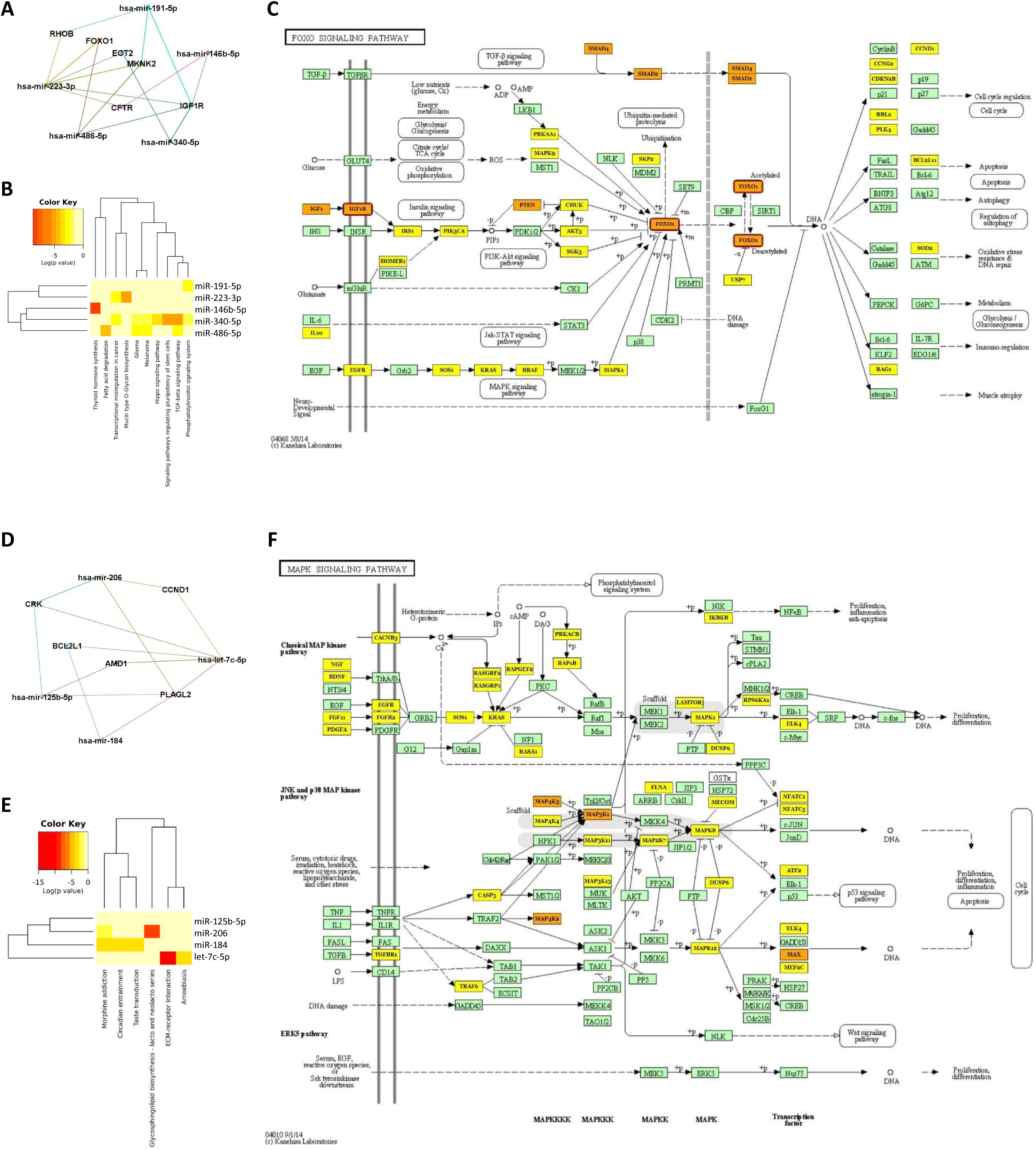
miRs responding to exercise in HT users may regulate the choice of energy source. **(A)** miR target analysis (miRNet) showing the minimal pathway map interconnecting miRs whose expression changed in response to exercise in EVs. **(B)** miR target pathway heatmap (miRPath v3) showing pathways that union between miRs that changed significantly in response to exercise in EVs. **(C)** Visualization of FOXO signaling pathway and key proteins regulated via significantly changed miRs in EVs in response to exercise. Green: no regulating miRs, Yellow: one regulating miR, Orange: > 1 regulation miR. **(D)** miR target analysis (miRNet) showing the minimal pathway map interconnecting miRs whose expression changed in response to exercise in HDL particles. **(E)** miR target pathway heatmap (miRPath v3) showing pathways that union between miRs that changed significantly in response to exercise in HDL particles. **(F)** Visualization of FOXO signaling pathway and key proteins regulated via significantly changed miRs in HDL particles in response to exercise. Green: no regulating miRs, Yellow: one regulating miR, Orange: > 1 regulation miR.

In HDL-carried miRs, the following interconnecting target proteins via miRNet were found: pleomorphic adenoma gene-like 2 (PLAGL2) was targeted by all four miRs; adapter molecule crk (CRK) was targeted by three miRs; B-cell lymphoma 2-like 1 (BCL2L1), adenosylmethionine decarboxylase 1 (AMD1), and cyclin D1 (CCND1) were targeted by different combinations of two miRs (Figure 4D). Further analysis via miRPath v3 with the KEGG signaling pathways union option showed six pathways to be targeted by the four miRs that had significant exercise response in HDLs (Figure 4E). When exploring genes’ union, we found seven KEGG signaling pathways that were regulated by those four miRs (Supplementary Table 16). Interestingly, several miRs that responded to exercise in HDL regulate target proteins in the MAPK signaling pathway (Figure 4F, Table S16). It is well established that MAPK signaling is required for various metabolic events (Gehart et al., 2010). Additionally, MAPK is a key regulator of gene transcription and metabolism in skeletal muscle in response to exercise and has been proven to promote fuel homeostasis (Kramer and Goodyear, 2007).

### Exercise increases the number of HDL particles in the blood

We observed, in both HT users and nonusers, an increase in APOA1 level and HDL particle number immediately after the VO_2peak_ test in a metabolomics analysis using serum (POST vs. PRE, Figures 5A-B). Our DB analysis of the plasma HDL fraction confirmed the increase in APOA1 when comparing POST vs. PRE in HT users, but there was no change observed in nonusers (Figure 5C). The DB analysis of the EV fraction showed a decrease in EV marker CD9 level in 1h POST vs. PRE comparison in nonusers, but this was not true in HT users (Figure 5D). The other EV marker, CD63, did not show exercise-induced differences (Figure 5E). The observed increase in HDL immediately after exercise is in agreement with previous studies showing similar results, that is, an increase in HDL particle number in response to exercise (Greene et al., 2012; Sondergaard et al., 2014). Because several methods of EV isolation also co-isolate lipoprotein particles, especially HDL due to similar density, it is important to note that not only the EV number but also the number of HDL particles may increase in response to acute exercise, potentially affecting the results.

**Figure 5.**
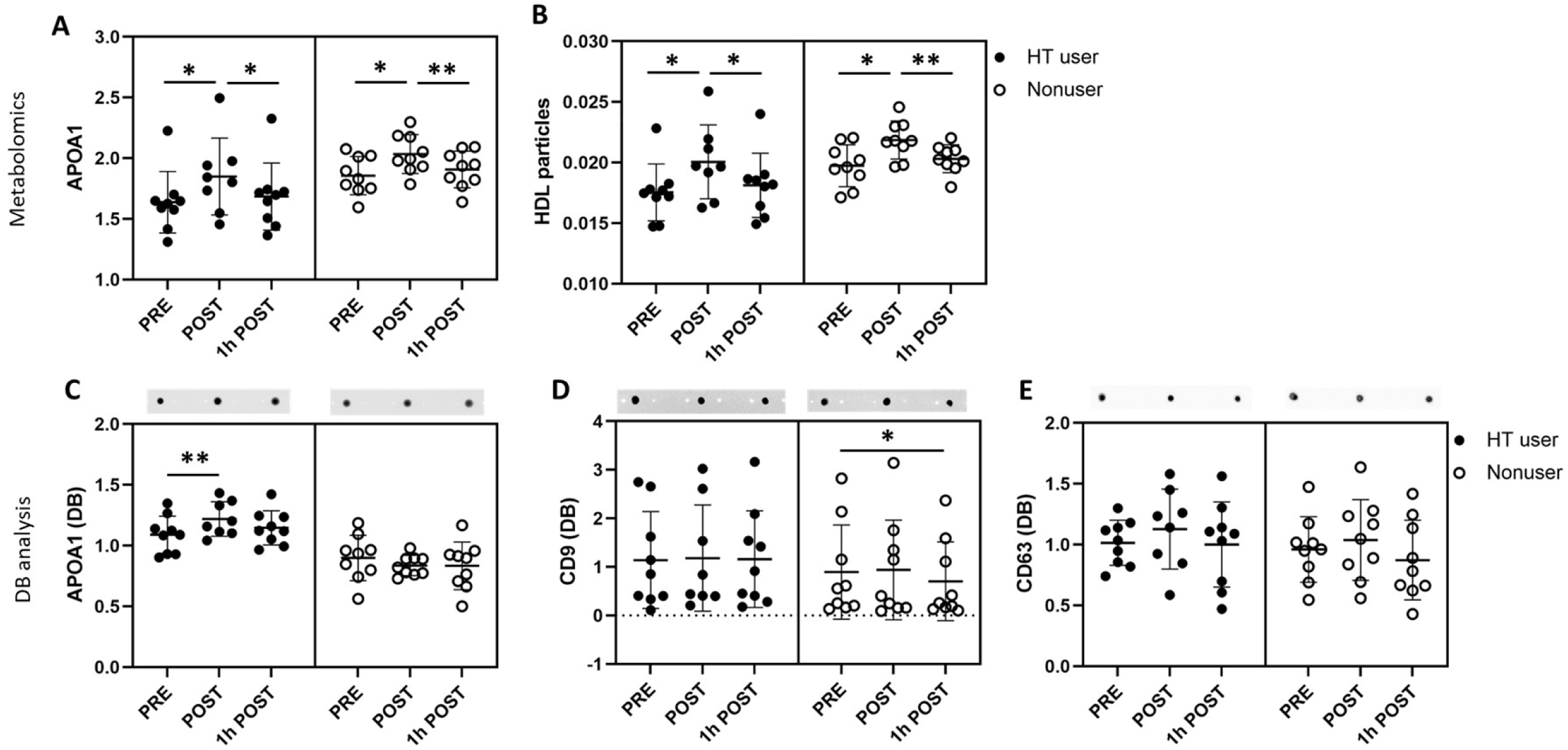
Exercise increases the number of HDL particles in the blood. **(A-B)** Metabolomic analysis of APOA1 and HDL particle number in serum. **(C-E)** DB analysis of APOA1, CD9 and CD63 level in isolated HDL (C) and EV (D-E) fractions.

## DISCUSSION

In the present study, we investigated sRNA molecules in two circulating signal transduction vehicles: EV and HDL particles. Our results revealed that EV and HDL particles have distinct sRNA content, with both carrying a substantial amount of nonhost sRNA. Of the host sRNAs, miRs were the most numerous RNA species present in EVs, and rDRs were most common in HDL particles, although the most abundant host sRNAs belonged to the miR class in both particles. We observed an exercise-induced increase in the relative miR abundance in EVs and the tDR abundance in HDL particles. By focusing on differentially expressed miRs, we demonstrated that the estrogen deficiency caused by menopause blunts the acute exercise-induced systemic miR response in both EV and HDL particles.

Excitingly, we show that nonhost sRNA represents a large share of the sRNA species in both carrier particles. Previously, Wang *et al*. (2012) observed that human plasma contains a significant amount (∼40%) of nonhost RNA (Wang et al., 2012). To further support our findings, Allen *et al*. (2018) showed that, in isolated mouse HDL, RNA of nonhost origin represents a larger relative share than host RNA (Allen et al., 2018). This was also true for our human HDL samples. In our study, the most common nonhost sRNA species in HDL were rDRs, which, similarly to Allen *et al*. (2018), were mostly mapped to a bacterial origin. In our data, the most abundant nonhost sRNA sequences were mapped to the human microbiome, suggesting an interconnection between the microbiome and sRNA-transporting particles. Supporting our findings, Roume *et al*. 2013 observed that samples derived from the human gastrointestinal tract were significantly enriched in sRNAs as compared to microbial samples derived from other environments (Roume et al., 2013). Accordingly, Wang *et al*. (2012) suggest that human gastrointestinal microbiota may actively synthesize sRNA molecules, which are then reflected in human blood (Wang et al., 2012). Our results support the hypothesis presented by Wang *et al*. (2012), which postulates that exogenous RNA found in plasma may mediate the human-microbiome interaction and further suggests that EV and HDL particles are signal transduction vehicles mediating these interactions.

In addition to less well characterized nonhost sRNAs, EV and HDL particles also carry a wide range of sRNAs mapped to the human genome. Our finding that EVs had, in general, a larger number of individual miR reads as compared with HDL particles supports the role of EVs as key mediators of miR-exchange between cells. Previously, Huang *et al*. (2013) reported a similar finding, showing that, in exosomes isolated from human plasma, miRs represent the largest RNA population (76.2% of all mappable reads) (Huang et al., 2013). EVs, in particular, are well-established miR carrier particles (Yanez-Mo et al., 2015), and miR release from EVs into recipient cells has been shown to induce various effects, such as regulating protein expression (Valadi et al., 2007). More recently, HDL particles have also been proven act as sRNA carriers that deliver miRs to recipient cells, such as hepatocytes (Vickers and Michell, 2021; Vickers et al., 2011).

Both EV and HDL particles have the potential to contribute to exercise adaptations by acting as exercise-responsive sRNA carrier particles. We observed several up-regulated rDR species in nonusers immediately after exercise in both carrier particles, even though the relative abundancy of rDRs decreased in both EV and HDL particles in response to exercise. Also, several rDR species were up-regulated 1h after exercise in the EVs of HT users. Strikingly, we noticed that all the exercise-responsive rDR species in EVs and the majority of those in HDL particles belonged to mitochondrial ribosomal subunits. Endurance exercise is well known to increase mitochondrial and ribosome biogenesis in skeletal muscle (Figueiredo and McCarthy, 2019; Mesquita et al., 2021), but recent research has shifted the perception of mitochondria from a simple energy-serving organelle to a key player controlling a variety of cellular signaling events (Kim, 2017). Indeed, mitochondrial peptides are now known as a class of circulating signaling molecules that regulate metabolism (Kim et al., 2017). For example, the mitochondrial peptide humanin, which originates from the mitochondrial 16S rRNA gene, has been linked to improved insulin action (Kuliawat et al., 2013). It was recently shown in humans that acute endurance exercise stimulates circulating levels of mitochondrial peptides (von Walden et al., 2021). We speculate that the observed changes in mitochondrial rDRs may reflect the response of muscle tissue to exercise, but the potential role of rDRs in coordinating metabolism remains to be uncovered.

Interestingly, we found an increase in the relative amount of tDRs in HDL in response to exercise. In contrast, several tDR species were downregulated in response to exercise in HDL particles in nonusers. Torres *et al*. (2021) recently proposed that extracellular tDRs may act as important paracrine signaling molecules whose activity depends on the mechanisms of the release and the capturing of recipient cells (Torres and Marti, 2021). To support this theory, the selective loading of tRNAs has been established in EVs (Chiou et al., 2018; Nolte-’t Hoen et al., 2012). Furthermore, previously, Lee *et al*. (2009) showed that tRNA-derived RNA fragments (i.e., tDRs) are not random by-products of tRNA degradation but, rather, an abundant and novel class of short RNAs with specific but, to a large extent, unknown biological roles (Lee et al., 2009). More recently, it was shown that, similar to miRs, tDRs are capable of down-regulating target genes via transcript cleavage *in vitro* (Li et al., 2012). According to the previous literature, we speculate that the observed changes in tDRs in HDL particles in response to exercise may serve gene-regulating roles, but more research is warranted to elucidate the potential effects on metabolism.

Previous studies by us (Karvinen et al., 2020) and others (Sapp et al., 2017; Silva et al., 2017) using various study populations have found c-miRs to respond to acute exercise, indicating that exercise normally triggers a detectable miR-response, underscoring the role of miRs as mediators of exercise adaptations. Strikingly, we observed that estrogen deficiency caused by menopause blunts the acute exercise-induced c-miR-response in both EV and HDL particles. This study showed that, immediately after the exercise test, the miRs 191-5p, 146b-5p, and 223-3p were upregulated, and 1h after exercise, miR-486-5p was upregulated, and miR-340-5p downregulated in the EVs of HT users, whereas we observed no response in nonusers. Of these miRs, miR-191-5p, 223-3p, and 486-5p were also among the twenty most highly expressed miRs, highlighting their role in coordinating physiological processes.

In contrast to our results, previously, Oliveira *et al*. found that miR-191-5p was down-regulated in response to acute exercise in rat serum (Oliveira et al., 2018). Several factors may have affected this observed difference. First, the exercise response between species may not be conserved. Second, Oliveira *et al*. used serum instead of plasma as a starting material, which causes the release of additional EVs during clot formation when preparing serum (Coumans et al., 2017; Wolf, 1967). Third, Oliveira *et al*. used a precipitation method to isolate EVs, which normally also includes protein-and lipoprotein-carried miRs, and thus, they may have examined signals from several different carrier particles. In cell studies, miR-191-5p has been shown to be involved in glucose metabolism via inhibiting the translocation of GLUT4 receptors, leading to decreased glucose uptake (Li et al., 2021; Yang et al., 2018). FOXO1, which is a target of the significantly up-regulated miRs 223-3p and 486-5p in our study, is a transcription factor regulated via the MAPK route. FOXO transcription factors have been demonstrated to be involved in angiogenesis, oxidative stress, and glucose metabolism (Matsumoto et al., 2007; Zhao et al., 2010). However, the current knowledge of the regulation of energy metabolism by miR-223-3p is contradictory. In cardiomyocytes, the overexpression of miR-223 has been reported to increase GLUT4 protein expression, leading to enhanced glucose uptake, while overexpression in human adipocytes has been associated with a decrease in GLUT4 protein content and diminished insulin-stimulated glucose uptake (Chuang et al., 2015; Lu et al., 2010). In turn, miR-486-5p has been shown to coordinate lipid metabolism and fatty acid degradation (Niculescu et al., 2018; Okamura et al., 2021). Furthermore, increased plasma levels of miR-486 have been correlated with dyslipidemia and hyperglycemia in patients with coronary artery disease (Simionescu et al., 2016), while the inhibition of miR-486 in a rodent model diminished plasma cholesterol and liver lipid content (Niculescu et al., 2018). Increased miR-146b-5p has been associated with decreased glucose metabolism and fatty-acid mobilization in obese subjects with nonalcoholic fatty liver disease (Latorre et al., 2017). Considering the literature, we speculate that the up-regulation of miRs 146b-5p, 223-3p, and 486-5p in HT users may play a role in exercise-induced energy substrate use adaptations. However, whether the c-miR response favors glucose or fatty acid oxidation may vary depending on the target tissue.

The function of HDL as a signal carrier is especially interesting because, after menopausal transition, both HDL-C and HDL particle numbers increase, but cardiovascular protection is diminished (Khoudary et al., 2016, 2018). In HDL particles, we observed that the miRs let-7c-5p, miR-125b-5p, miR-206, and miR-184 were upregulated both immediately and 1h after exercise in HT users, with no response observed in nonusers. Previous studies have found miR-125b expression to be increased in response to estrogen *in vitro* and *in vivo*, as well as providing evidence of fatty acid synthase as a functional target of miR-125b (Zhang et al., 2015b). Increased miR-125b has proven to inhibit lipid accumulation in hepatocytes due to decreased fatty acid uptake and synthesis and decreased triglyceride synthesis (Zhang et al., 2015b). On the basis of our study, miR-125b-5p is estrogen responsive in the context of exercise, but only in HDL particles.

Excitingly, two muscle-specific miRs (myomiR) (Horak et al., 2016), miR-486-5p in EVs and miR-206 in HDL particles, were increased in response to exercise. This finding indicates that, in addition to EVs, HDL particles also exchange miRs with muscle cells during exercise. Previously, HDL has been proven to transport and deliver functional miRs to cultured hepatocytes and endothelial cells (Tabet et al., 2014; Vickers et al., 2011). Also, miR-206 levels have been shown to increase in plasma in response to heavy endurance exercise, such as marathon running (Mooren et al., 2014). Due to the role of myomiRs (miR-1, miR-133a/b, miR-206, miR-208a/b, miR-486, and miR-499) in skeletal and/or cardiac muscle development, the up-regulation of these miRs in response to exercise is proposed to be relevant to exercise-induced adaptations to improve aerobic capacity (Mooren et al., 2014). Interestingly, miR-206 in the liver has been shown to prevent excess fat accumulation and hyperglycemia (Wu et al., 2017). Because HDL transports cholesterol to the liver and may simultaneously exchange other molecules with hepatocytes, we speculate that the miRs 125b-5p and 206 share a role in selecting an energy source in HT users during exercise, together with the EV-carried miRs 223-3p and 486-5p, possibly by decreasing fat uptake into the liver.

In the present study, only HT users exhibited an miR-response to exercise. Hence, we propose that c-miRs could provide a means of transmitting estrogen action during exercise, potentially by affecting mitochondrial function. The fuel selection of a contracting skeletal muscle depends on several factors, which originate at the mitochondrial level (Kuzmiak-Glancy and Willis, 2014; Overmyer et al., 2015). We have previously shown that estradiol (E_2_), which is the physiologically predominant estrogen, improves bioenergetic function in skeletal muscle by affecting mitochondrial membrane viscosity (Torres et al., 2018). Thus far, several miRs that translocate into the mitochondria have proven to play an important role in mitochondrial diseases, as well as controlling metabolic homeostasis (Sekar et al., 2020). Indeed, FOXO transcription factors act in crosstalk between mitochondria and the nucleus (Kim and Koh, 2017). In rodents and cell models, FOXO acts as a switch to lipid utilization as the major fuel substrate in skeletal muscle (Bastie et al., 2005; Furuyama et al., 2003). However, in humans, the effects of exercise on FOXO1 in skeletal muscle remain controversial (Tsintzas et al., 2006). The MAPK signaling cascade, in turn, is a chain of proteins that transports a signal from a receptor on the surface of the cell into the cytoplasm, regulating several pathways (Gehart et al., 2010). Previous studies have indicated that MAPKs directly interact with and translocate into mitochondria (Ballard-Croft et al., 2005). The estrogen-responsive miR-125b-5p, which was upregulated in HDL in response to exercise in HT users, may play an important role in modulating oxygen consumption and mitochondrial gene expression in adipocytes (Giroud et al., 2016). We speculate that c-miRs may function as a link between estrogen status and mitochondrial function, coordinating the metabolism in response to an acute bout of exercise. Interestingly, various exercise responses were seen in HDL particles, which may explain some controversies regarding the known associations between high HDL and high physical activity, aerobic capacity and reduced risk of cardio-metabolic diseases by shifting the focus from HDL cholesterol concentration to other functional properties of HDL particles (Kujala et al., 2013, 2019; Lehti et al., 2013; Vickers and Michell, 2021).

In summary, we are the first to show that EVs and HDL particles display distinct sRNA content, which is rich in nonhost RNA species. Furthermore, we show that only HT users exhibit an acute exercise-induced miR-response in two systemic carriers, that is, EV and HDL particles. In line with our previous observation that menopause leads to the diminished cardiovascular protection of HDL particles, estrogens also alter the miR-cargo of HDL particles in response to exercise. The exercise-induced c-miR response is regulated by estrogens and may contribute to fuel selection, potentially by affecting mitochondrial function in postmenopausal women.

### Study strengths and limitations

This study has several important strengths. First, it meticulously verified an isolation protocol enabling the separation of EV and HDL fractions before RNA isolation. Second, it relied on a combination of bleeding diaries and serum FSH levels rather than on self-report alone to accurately classify women as postmenopausal. Second, matching the groups for age, VO_2peak_, BMI, and body fat percentage allowed us to eliminate several confounding factors known to affect exercise response and lipid metabolism. Our study also has certain limitations. One is the relatively small number of participants, which is due to the cancellation of participant recruitment due to the COVID-19 pandemic and practical issues limiting the time scale for data collection. Thus, only nine HT users and matched nonusers could be recruited within the criteria stated previously. We noted that the APOA1 antibody also showed a positive signal in EV fractions of samples, suggesting the potential presence of HDL particles in the sample. It is, however, important to note that, according to recent findings, APOA1 may be locate in the EV corona (Toth et al., 2021). Nevertheless, total protein, WB, DB, EM, and IEM analysis confirmed that HDL particles were enriched in the HDL fraction. Previous studies have shown an increase in EV number in the circulation after a single bout of exercise (Brahmer et al., 2019; Fruhbeis et al., 2015; Oliveira et al., 2018; Whitham et al., 2018). Due to the limited volume of plasma samples and the methodology available, we were not able to measure the total number of EVs with the best suitable methods, such as flow cytometry. Nevertheless, we are confident that the results are relevant regardless of whether the changes observed are partly due to the increased number of EVs because only a few specific miRs showed a clear response to exercise stimulus in HT users, supporting highly coordinated regulatory mechanisms for miR packaging into EVs. In this study, we were not able to determine the origin or target tissue of EVs or HDL particles, and thus, the potential effects of the significantly altered miRs may vary depending on the receiving tissue. Also, we were not able to control for blood volume changes during and after the acute exercise test, but it is unlikely that plasma volume changes explain our findings given our study design.

## ACKNOWLEDGEMENTS

This study was funded by grants from the Academy of Finland (grant numbers 332946 to SK and 309504, 314181 and 335249 to EKL). We would like to thank the laboratory staff at the Faculty of Sport and Health Sciences for their invaluable assistance in the data collection. We also want to thank the women who participated in the EsmiRs study for their time and effort.

## AUTHOR CONTRIBUTIONS

SK performed the sample analysis, ran the corresponding statistical analysis, and prepared the first version of the manuscript. TMK prepared the NGS libraries under the supervision of TS. TS and TMK performed NGS and conducted the preliminary processing of the raw data. TMK installed the TIGER pipeline and analyzed the NGS results under the supervision of EKL. JEK and HKJ performed the VO_2peak_ tests and harvested the blood samples. VJ performed ultracentrifugation and qPCR analysis for a subset of the samples under the supervision of SK. JAL performed clinical analysis of the study subjects. ML supervised the HDL isolation protocol. EKL designed the study, offered funding and conceived and supervised the study. UKM critically contributed to the design and implementation of the study. SK and EKL wrote the final version of the manuscript. All authors critically commented on the manuscript draft and approved the final version of the manuscript.

## METHODS

The present study investigated plasma samples from a subset of the “Estrogen, microRNAs and the risk of metabolic dysfunction (EsmiRs) study” (Laakkonen et al., 2021) consisting of 18 postmenopausal women (age 52–57 years, Supplementary Table 1). Of these women, nine used estrogen-based hormonal therapy (HT), and nine were nonusers. We carried out an NGS analysis with 14 purified EV and HDL samples from HT users and nonusers (n = 7/group). Study groups were matched for age, peak aerobic capacity (VO_2peak_), body mass index (BMI), and body fat percentage.

The study was approved by the Ethics Committee of the Central Finland Health Care District (KSSHP Dnro 9U/2018). Written informed consent was obtained from all participants before the study. The study was conducted in conformity with the Declaration of Helsinki.

### Antropometrics

The body mass index (BMI) of the study subjects was calculated by dividing body mass (kg) by height squared (m^2^). Body fat percentage (BF%) was measured via dual-energy X-ray absorptiometry scans (LUNAR Prodigy; GE Healthcare, Chicago, IL) after overnight fasting.

### 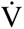O_2peak_ test

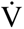O_2peak_ was measured with a Vmax Encore 92 metabolic cart (Sensormedics, Yorba Linda, CA, USA) during a maximal incremental bicycle ergometer (Ergoselect 200, Ergoline GmbH, Bitz, Germany) test. Testing was performed after overnight fasting, and study subjects were instructed to abstain from alcohol intake, as well as strenuous exercise, for 48-hours before the test. The 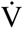O_2peak_ test comprised both submaximal and maximal phases. First, study subjects cycled for 4 min at an intensity of 20 watts (*W*), after which the workload was increased by 20 *W* every 4 min until a respiratory exchange ratio of 1.0 was reached. Thereafter, the intensity was increased by 1 *W*/3 s to a total of 20 *W*/min. The test continued until volitional exhaustion, after which the participants performed a 5-minute cooldown at an intensity of 50 *W*. Gas exchange was recorded as 10 s rolling averages, and the highest continuous 30-s 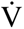O_2_ period was selected to represent participants’ absolute 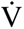O_2peak_. Respiratory gas exchange measurement was unreliable in two HT users due to metabolic cart failure or mask-wearing difficulties. For these two participants, 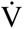O_2peak_ was determined based on maximal workload, using the equation created by Storer *et al*. (1990). 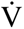O_2peak_ was also scaled relative to body weight (ml/kg/min) and lean body mass (ml/kg LBM/min).

### Leisure-time physical activity (LTPA)

Leisure-time physical activity (LTPA) was assessed from both self-reported and measured data. Self-reported LTPA activity was assessed using a questionnaire (Kujala et al., 1998). Briefly, LTPA was assessed based on three questions describing the monthly frequency, mean duration and mean intensity of LTPA to elicit participants’ opinions regarding their overall LTPA levels. Metabolic equivalent (MET) values were assigned for each activity. Using these, the total volume of LTPA was calculated as a product of duration, intensity, and frequency. The total LTPA was described as MET hours per day.

Measured LTPA was assessed using GT3X+ and wGT3X+ ActiGraph accelerometers (Pensacola, Florida, USA), as described previously (Laakkonen et al., 2017). Briefly, accelerometers were placed on the right hip and used for 7 consecutive days during waking hours, excluding bathing and other water-based activities. In addition, participants were asked to record their wake-up times and working hours in a diary. Raw data from the accelerometers were collected at 60 Hz and filtered, after which they were converted into 60-s epoch counts. Further data analysis was performed by using a customized Excel-based program, as described previously (Laakkonen et al., 2017). Entries about working hours in activity diaries were used to distinguish LTPA from total daily PA. Data normalization was performed for a 10-hour period of being awake. Activity was described as total counts for 10-hour LTPA.

### Blood sampling and analysis

To measure the blood lipid and hormone levels of the participants, blood samples were harvested after overnight fasting between 7:00 and 10:00 a.m. Blood was drawn from the antecubital vein in a supine position. To separate the serum, whole blood was left to clot for 30 minutes at room temperature, after which it was centrifuged at 2,200 × g, and the sera were aliquoted and stored at – 80 °C, as described previously (Kovanen et al., 2018). Then, TC, HDL-C, LDL-C, and TGs were determined using a KONELAB 20 XTi analyzer (Thermo Fischer Scientific, Finland) according to the manufacturer’s instructions, and E_2_ and FSH levels were measured from the serum using an IMMULITE® 2000 XPi (Siemens Healthcare Diagnostics, UK), following the manufacturer’s instructions. Serum HDL particle numbers and APOA1 were analyzed using a targeted high-throughput nuclear magnetic resonance (NMR) spectroscopy platform (Nightingale Health Ltd., Helsinki, Finland; biomarker quantification v. 2020).

During the VO_2peak_ test, blood samples were harvested after overnight fasting at 7:15 a.m. A total of 18 ml of peripheral blood was collected into two 9 ml Vacuette® EDTA K3 tubes at three time points: PRE, POST and 1 h POST exercise test. The whole blood was mixed with 18 ml of RPMI-medium, and white blood cells from plasma were isolated with Ficoll-Paque™ PLUS medium (GE healthcare). Thereafter, the plasma was stored at -80°C before being used for HDL and EV isolation.

### EV and HDL isolation

The fraction containing HDL and EVs was isolated from plasma samples harvested during the VO_2peak_ test using sequential ultracentrifugation with potassium bromide (KBr). First, plasma samples were defrosted overnight, density-adjusted to 1.063 g/ml, and ultracentrifuged (60,000 rpm, 4 °C, Beckman, 70 Ti rotor) for 20 hours. To purify the HDL and EV fraction, 5 ml of solution was collected from the bottom of each tube and the density readjusted to 1.21 g/ml. The density-adjusted solution was centrifuged (60,000 rpm, 4 °C) for 48 hours. The HDL and EV containing fraction (5 ml) was collected from the top of each tube. Prior to SEC, 4 ml of each sample were concentrated with Amicon ultra Centrifugal filters (Cat no. UFC510024, Merck). Samples were centrifuged (14 000 × g, 4 °C, Heraeus Fresco 17 centrifuge) for 10 minutes to concentrate the HDL and EV fraction. The concentrated HDL and EV fraction was further centrifuged (1000 × g, 4 °C) for 2 minutes to obtain 60 µl of concentrate.

HDL was separated from EVs using SEC (IZON qEVoriginal, 35 nm, SP5). Concentrated samples (60 µl) were taken to room temperature to thaw on ice. The SEC column was taken to room temperature, positioned in an upright position, and used according to the manufacturer’s instructions. Then, 440 µl filtered (0.2 µm) PBS (pH 7.4) was added to the thawed samples in order to reach the 500 µl sample volume recommended for SEC. Each sample (500 µl) was loaded into the column, and the void volume (3 ml) was discarded. Thereafter, EV and HDL zone volumes were collected into sixteen 0.5 ml fractions. The EV zone volume was collected into fractions 1–3, as informed by the manufacturer, and verified via DB, EM, and IEM analysis (see “Validating EV and HDL isolation”). Based on total protein and WB, DB, and EM analysis, the majority of HDL was located between fractions 7 and 11 (see “Validating EV and HDL isolation”). EVs and HDL were concentrated separately from these fractions, as described above, to reduce the volume for RNA extraction.

### Validating EV and HDL isolation

The successful isolation of HDL particles and EVs via SEC was investigated by measuring total protein concentration and utilizing WB, DB, EM, and IEM. For validation purposes, representative HDL+EV samples isolated from serum were used.

First, to determine the zone of HDL, total protein was measured from SEC fractions using a bicinchoninic acid protein assay (Pierce Biotechnology, Rockford, IL) with an automated KoneLab instrument (Thermo Scientific, Vantaa, Finland). Based on total protein, the WB and EM majority of HDL was verified as located between fractions 7 and 11 (Figures 1A-F). HDL was concentrated from these fractions, as described above, to reduce the volume for RNA extraction. Second, WB was carried out for 15 µl of the same fractions using APOA1 antibody to verify the presence of HDL particles in the most protein-rich fractions (Figure 1B). For WB, samples were solubilized in Laemmli sample buffer and heated at 95°C for 10 min to denature proteins and separated via SDS-PAGE for 40–60 min at 270 V using 4–20% stain-free gradient gels on Criterion electrophoresis cell (Bio-Rad Laboratories, Richmond, CA). Thereafter, the proteins were transferred to nitrocellulose membranes in a Turbo blotter (Trans-Blot Turbo Blotting System, 170-4155, Bio-Rad Laboratories). The total protein amount was visualized using the stain-free image of the blot (Figure 1B). The membrane was blocked in commercial blocking buffer (Odysseu Blocking Buffer [PBS], Licor) for 2 h and then incubated overnight at 4°C with primary antibody to measure APOA1 (1:10 000, cat# ab52945, RRID:AB_2056661, Abcam, Cambridge, United Kingdom). Afterward, the primary antibody incubation membrane was washed in TBS-T, incubated with suitable secondary antibody (1: 10 000), and diluted in 1:1 Pierce blocking buffer and TBS-T for 1 h, followed by washing in TBS-T. Proteins were visualized via fluorescence using ChemiDoc XRS, in combination with Quantity One software (Version 4.6.3. Bio-Rad Laboratories). Based on both total protein measurement and WB images of APOA1, the majority of HDL was located between fractions 7 and 11 (Figure 1A-B).

Because the protein level of EV samples was not detectable via traditional WB (*data not shown*), DB analysis were carried out for all sixteen SEC fractions similarly to WB, except that the 1 µl of sample was blotted straight onto nitrocellulose membrane (Figure 1C) and total protein was visualized using PonceauS staining. Visualizing APOA1 content was performed as described above, and in addition, the levels of CD9 (1:500; cat# ab92726, RRID:AB_10561589, Abcam) and CD63 (1:500; cat# ab59479, RRID:AB_940915, Abcam) were visualized. The BD analysis of the SEC fractions confirmed the location of EVs in fractions 1–3 and the enrichment of HDL in fractions 7–11 (Figure 1C).

For further DB, EM, and IEM analysis, four representative serum samples containing EV and HDL particles were first isolated via density-adjusted sequential ultracentrifugation, as described in the “EV and HDL isolation” section. Due to the high protein concentration in the HDL fraction as compared with the EV+HDL and EV fractions, a dilution series of the HDL fraction was run to determine the suitable dilution of HDL for optimal performance for the antibodies used (Figure 1D). An undiluted EV sample was used as a positive control for EV antibodies. According to our findings, the HDL sample that was diluted to 1:100 in filtered PBS (pH 7.4), when visualized together with EV+HDL and EV samples, was optimal for the detection of antibodies (Figure 1D-E).

Samples containing EVs and HDL (the fraction collected after sequential ultracentrifugation) and EVs and HDL separately (the fractions after SEC) were prepared for EM and IEM analysis, as described previously (Karvinen et al., 2020). For analysis, EV+HDL, EV, and HDL samples were loaded on 200 mesh grids, fixed with 2% paraformaldehyde solution (PFA), stained with 2% neutral uranyl acetate, and embedded in a uranyl acetate and methyl cellulose mixture (1.8/0.4%). Samples were viewed with transmission EM using a Jeol JEM-1400 (Jeol Ltd., Tokyo, Japan) operating at 80 kV. Images were taken with a Gatan Orius SC 1000B CCD-camera (Gatan Inc., United States) (Puhka et al., 2017).

The IEM staining was performed similarly to EM, with the addition of an immunostaining step. Briefly, after being loaded onto 200 mesh copper grids, the samples were blocked with 0.5% BSA in 0.1 M NaPO4 buffer (pH 7.0) and incubated with anti-CD9 (1:100 dilution, BioSite MEM-61) in 0.1% BSA/NaPO4 buffer. Thereafter, the samples were incubated with 10 nm gold-conjugated goat anti-mouse IgG (1:80 dilution, BBI Solutions, Cardiff, UK) in 0.1% BSA/NaPO4 buffer, washed with the NaPO4 buffer and deionized water, negatively stained with 2% neutral uranyl acetate, and embedded in methyl cellulose uranyl acetate mixture (1.8/0.4 %) (Figure 1F).

### RNA extraction

RNeasy Serum/Plasma Kit (217184, Qiagen) was used to isolate the RNA from the EVs and HDL particles according to the manufacturer’s instructions. Before isolation, 140 µl of filtered (0.2 µm) PBS (pH 7.4) was used to obtain the recommended starting volume of 200µl. Spike-in cel-miR-39-3p (MS00019789, Qiagen) was used as an internal control.

### sRNA library preparation and NGS

sRNA Library preparation was carried out utilizing QIAseq miRNA Library Preparation Kit (Qiagen, Germany, Cat no. 331505) according to manufacturer’s instructions, using multiplexing adapters. Briefly, sRNA fractions were first ligated to adapters from both the 5’ and 3’ ends, reverse transcribed into cDNA using UMI-assigning primers, and purified using magnetic beads. A universal indexing sequence to distinguish individual samples was added to each sample in the reverse transcription step. The sRNA libraries were then amplified with PCR (Eppendorf, Germany), purified, and eluted into 18 ul of nuclease-free water. The sRNA libraries were stored at -20 C until further analysis. A Qubit fluorometer (Invitrogen, USA) was used to measure library concentrations, and the libraries were subsequently diluted and pooled into an equimolar mixture containing 1.8 pM per sample prior sequencing. The sequencing of the sRNA libraries was carried out with NextSeq 500 (Illumina, USA), using a NextSeq 500/550 High Output Kit v. 2.5 with 75 cycles (Illumina, USA) to produce 75-base pair single-end reads.

### Raw sRNA data processing and alignment

Illumina sequencing output data were converted to .fastq data format using bcl2fastq software (v.2.20, Illumina, USA). Quality controls were performed throughout the analysis with FastQC (Andrews S. [2010] FastQC: A Quality Control Tool for High Throughput Sequence Data. Available online at: http://www.bioinformatics.babraham.ac.uk/projects/fastqc/). The QIAseq sequencing adapters were trimmed from the 3’ end of the reads (5’ –AACTGTAGGCACCATCAAT-3’) with the FastX-Toolkit (Hannon GJ. (2010) FASTX-Toolkit. http://hannonlab.cshl.edu/fastx_toolkit), using default parameters with a minimum alignment length of -M 19 and a minimum read length requirement of -l 20. Only clipped reads were selected for downstream analysis. For the analysis of miRs (but not rRNAs or tRNAs), the reads were trimmed to 22 bp using a FastX-Toolkit to enrich miR-sequences. Finally, the reads were quality-filtered with a FastX-Toolkit using the parameters *-q 25* and *-p 90*, meaning that 90% of the reads should have an average Phred score of 25. Only high-quality reads were selected for alignment to a reference genome. Before alignment, all four sample lanes were merged to obtain the overall sample read count and ensure better-quality mapping. Samples that had <600,000 raw reads were excluded from the analyses. Alignment was done using Bowtie (Langmead et al., 2009). Default parameters were utilized for reported alignments so that only one best alignment for a read was used as output, even if there were multiple possible alignments. miRs were aligned to miR base version 22 (Kozomara et al., 2019), rDRs were aligned to the full set of human rRNA (downloaded from RNAcentral in January 2022), and tDRs were aligned to the high-confidence tRNA gene set from GtRNAdb (Chan and Lowe, 2016).

### TIGER pipeline analysis of sRNA

We examined the sRNA species in EV and HDL particles by utilizing a novel data analysis pipeline entitled “Tools for Integrative Genome analysis of Extracellular sRNAs (TIGER)” (Allen et al., 2018). This TIGER analysis uses genome and database alignments to categorize sRNA sequences based on their origin. It also outputs the most abundant hundred sequences in each sample. The TIGER pipeline was used to produce the results presented in Figures 1G–1L, as well as Figures 2A–B.

### Differential expression analysis of sRNAs

The DE analyses of c-miR counts were performed using DESeq2. Briefly, a DE analysis was run for n = 7/group with the following six samples from each study subject: EV PRE, EV POST, EV, 1h POST, HDL PRE, HDL POST, and HDL 1h POST. Samples with <500 total miR counts were excluded from the analysis (one sample in addition to the two samples excluded earlier due to low raw reads). Low-count miRs were filtered out. In order to be included in the DE analysis, an miR had to have counts in 70% of the samples of HT users or nonusers or in 5/7 samples in a subgroup (time point PRE, POST, or 1h POST). The reads were normalized with DESeq’s median of ratios normalization method (Love et al., 2014) . To compare each individual’s samples at different time points, paired analysis was performed by including the test subject as a factor in the design formula. c-miRs that had false discovery rate (FDR) <0.05 were considered differentially expressed. Normalized miRs were examined at each time point in EV and HDL particles in Figure 2C, and DE analysis results are shown in Figures 3A-D. The original and normalized miR counts are presented in Supplementary Figure 1. The DE analyses of rDRs and tDRs were conducted using essentially the same methods as with miRs. However, because there were more samples with low read counts than with miRs (two for rDRs and four for tDRs), we decided to replace those samples with group averages.

### miR target analysis and signaling pathway heatmap

After finding significantly altered miRs after exercise in EVs and HDL particles, we performed a miR target analysis via miRNet (https://www.mirnet.ca/). For EV miR analysis, the tissue was set to exosomes, and for miRs carried in HDL particles, the tissue was unspecified. For all analysis performed in miRNet, minimal pathway analysis was used. To investigate the miR–mRNA signaling pathway interaction, we used mirPath v.3. We used the Human database and the microT-CDS for mRNA target prediction, with default threshold values (p ≤0.05, MicroT threshold 0.8) and FDR correction.

### cDNA synthesis and qPCR

After RNA isolation, cDNA synthesis was performed with an miScript II RT Kit (218161, Qiagen) using HiFlex buffer according to manufacturer’s instructions. cDNA synthesis was performed using 12 µl of non-diluted template RNA, and PCR protocol was carried out in a standard PCR device (Eppendorf AG 22331, Hamburg). For the qPCR runs, 1 μl of non-diluted cDNA was used per well, and samples were run as duplicates. We studied miRs 191-5p, -223-3p, and -486 from the EV fraction and let-7c-5p and miR-125b-5p from the HDL fraction. The sequence of the universal primer was 5’-GAATCGAGCACCAGTTACGC-3’. The sequences of miRNA-specific primers were obtained from miRBase and were as follows:

hsa-let-7c-5p, MIMAT0000064: 5’-TGAGGTAGTAGGTTGTATGGTT-3’,
hsa-mir-125b-5p, MIMAT0000423: 5’-TCCCTGAGACCCTAACTTGTGA -3’,
hsa-mir-191-5p, MIMAT0000440: 5’-CAACGGAATCCCAAAAGCAGCTG -3’
hsa-mir-223-3p, MIMAT0000280: 5’-TGTCAGTTTGTCAAATACCCCA-3’
hsa-mir-486-5p, MIMAT0002177: 5’-TCCTGTACTGAGCTGCCCCGAG-3’

The qPCR protocol was as follows: 95°C (15 min, activation), 94°C (15 s), 55°C (30 s), and 70°C (30 s) with 40 cycles (CFX384™ Real-Time PCR Detection System, Bio-Rad). The relative expression of the studied miRs was calculated using equation 2^−Δ *Cq*^, where ΔCq = target miR Cq-value/housekeeping Cq-value and was expressed as the fold change to corresponding (HT user or nonuser) PRE values.

### Statistics

Statistical analyses were performed using IBM SPSS for Windows 24 statistical software.

Data are presented using means and standard deviations. To determine the normality of variables, the Shapiro-Wilk test was applied, and the skewness and kurtosis of the distributions of variables were interpreted. For the study subject characteristics (Table S1), statistical analysis was run via a Student’s *t*-test for the parameters that were normally distributed and a Mann-Whitney U-test for the parameters that did not meet normal distribution criteria. When examining pairwise differences between time points, a paired-samples *t*-test was used for normally distributed data, and a Wilcoxon signed-rank test was used for the data that did not meet normal distribution criteria. In all analyses, a p-value < 0.05 was considered to indicate statistical significance.

---

**Figure S1.**
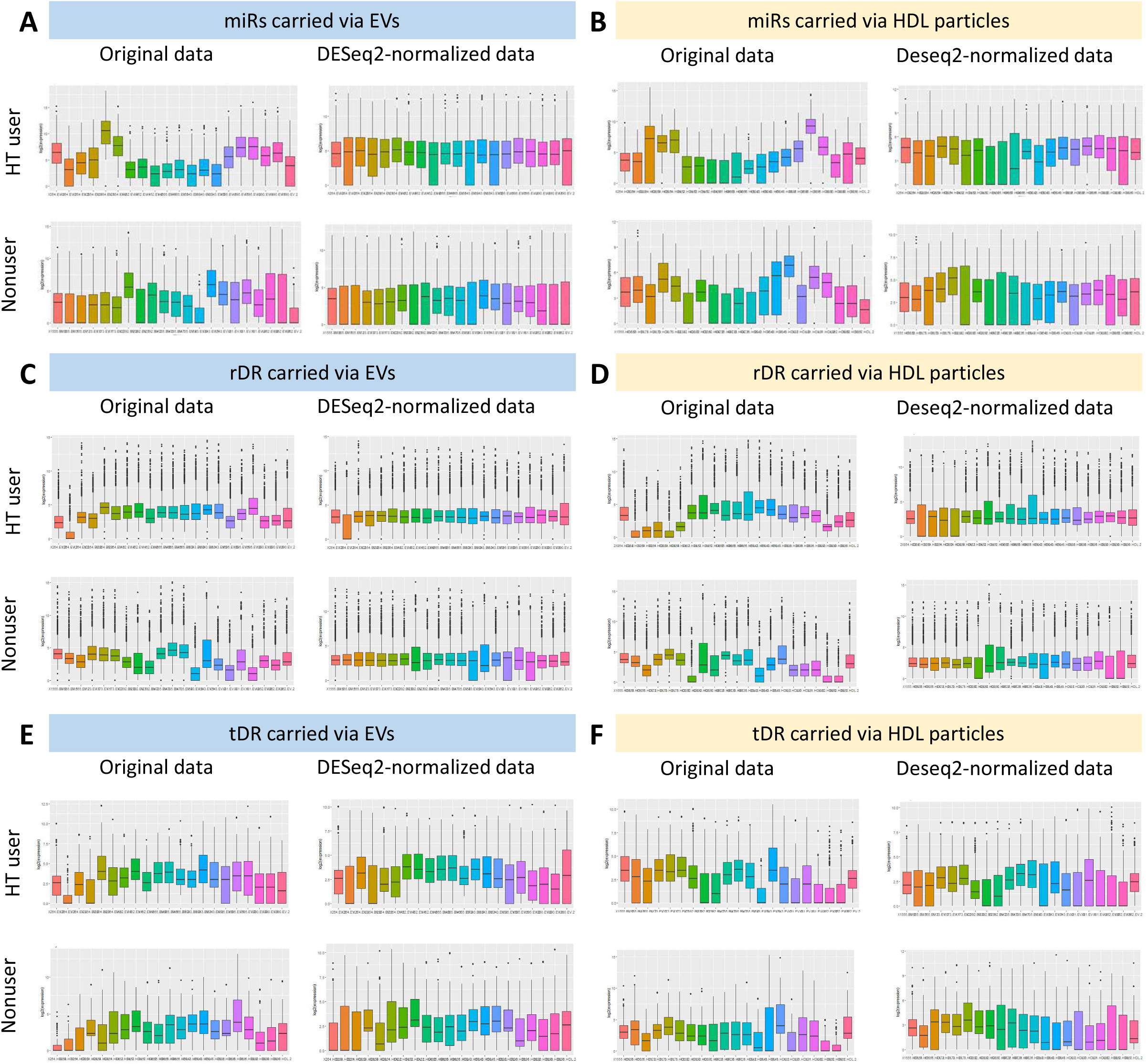

**Figure S2.**
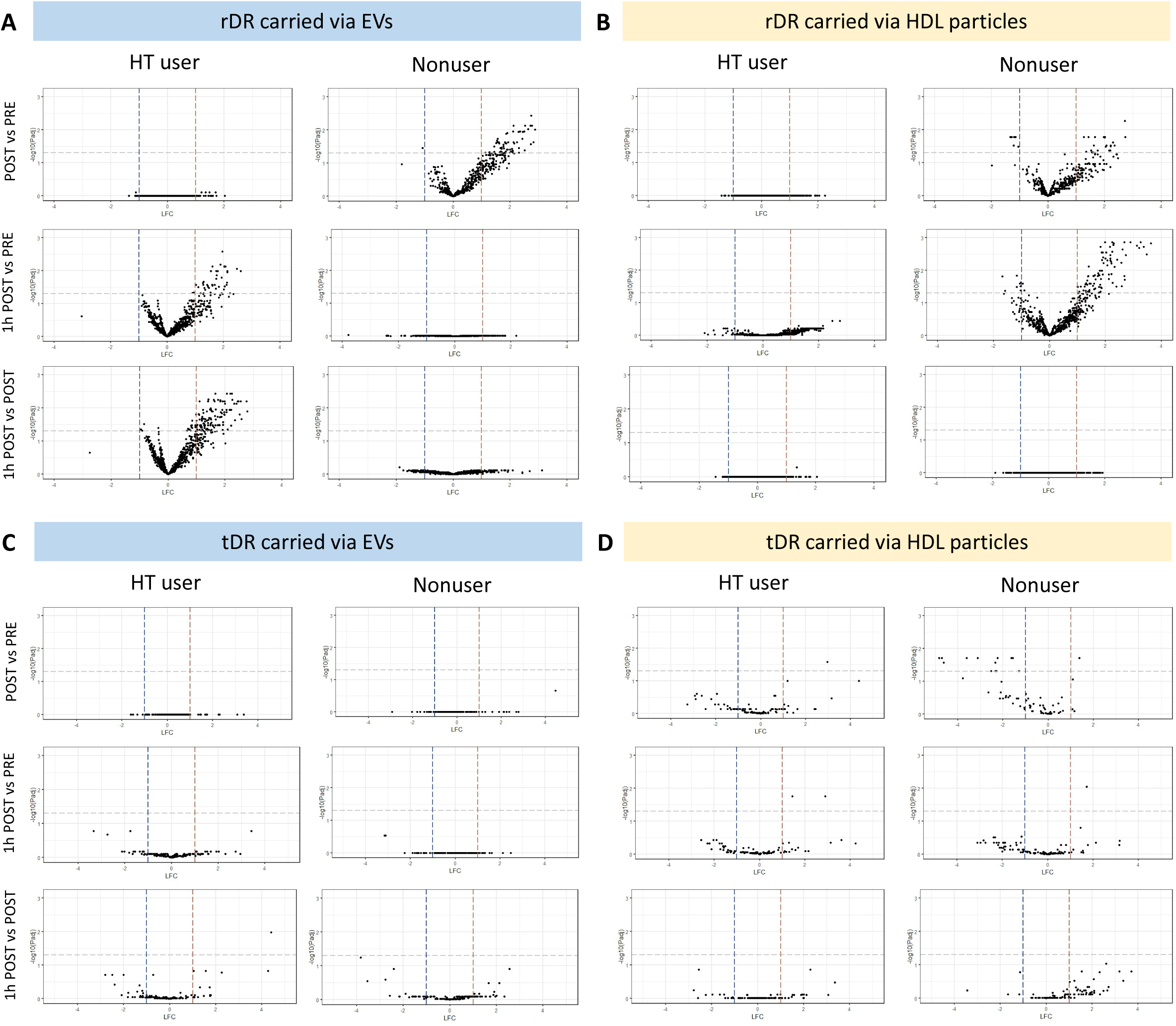

