## Supplemental information for "Extracellular vesicles and high-density lipoproteins: Exercise and estrogen-responsive small RNA carriers"

### Supplemental figure legends

**Figure S1. Original and DESeq2-normalized miRs, rDR, and tDR.**
**(A-B)** Visualization of original and DESeq2-normalized data of miRs carried in EVs and HDL particles in HT users and nonusers.
**(C-D)** Visualization of original and DESeq2-normalized data on rDRs carried in EVs and HDL particles in HT users and nonusers.
**(E-F)** Visualization of original and DESeq2-normalized data on tDRs carried in EVs and HDL particles in HT users and nonusers.

**Figure S2. HT users and nonuser have a distinct response to acute exercise in rDR and tDR carried via EV and HDL particles.**
**(A-B)** Volcano plots showing differential expression of rDRs between time points in EV and HDL particles in HT users and nonusers.

**(C-D)** Volcano plots showing differential expression of tDRs between time points in EV and HDL particles in HT users and nonusers.

**Table S1**: Characteristics of all subjects (n = 9) and subjects used in NGS analysis (n = 7).

| **Variable** | **HT user (n = 9)** | **nonuser (n = 9)** | **p-value** | **HT user (n = 7)** | **nonuser (n = 7)** | **p-value** |
|---|---|---|---|---|---|---|
| ***Background characteristics*** | | | | | | |
| Age (y)a | 55.45±0.46 | 56.10±0.52 | 0.364 | 55.26±1.52 | 55.65±1.43 | 0.633 |
| BMI (kg/m2)b | 24.82±1.08 | 24.58±0.83 | 0.566 | 23.67±2.56 | 24.13±2.58 | 0.180 |
| Body fat (%)a | 38.92±1.46 | 37.65±1.80 | 0.592 | 37.87±4.42 | 36.92±5.78 | 0.920 |
| Lean body mass (kg)a | 40.21±3.73 | 41.51±2.91 | 0.419 | 38.58±2.17 | 42.23±2.90 | **0.021** |
| VO2peak (ml/kg/min)a | 26.97±1.33 | 27.31±1.38 | 0.860 | 26.86±4.45 | 27.74±4.50 | 0.720 |
| VO2peak (ml/kg of lean body mass/min)a | 46.57±6.67 | 47.93±5.53 | 0.643 | 45.69±7.14 | 48.35±5.09 | 0.438 |
| Self-reported LTPA (MET hours/day)b | 4.52±4.28 | 3.84±3.70 | 0.757 | 4.22±4.30 | 4.27±4.16 | 0.949 |
| Measured LTPA (total counts/day)a | 4.09*10^5±6.36*10^4 | 5.61*10^5±1.80*10^4 | **0.046** | 4.25*10^5±2.10*10^5 | 5.52*10^5±5.91*10^4 | 0.166 |
| ***Blood variables*** | | | | | | |
| HDL-C (mmol/l)a | 1.76±0.19 | 2.22±0.13 | 0.057 | 1.91±0.50 | 2.30±0.39 | 0.128 |
| LDL-C (mmol/l)a | 3.22±0.24 | 3.53±0.46 | 0.551 | 3.04±0.70 | 3.07±0.83 | 0.943 |
| TC (mmol/l)b | 5.39±0.18 | 6.24±0.45 | 0.102 | 5.36±0.59 | 5.85±0.76 | 0.204 |
| TG (mmol/l)a | 1.12±0.13 | 1.31±0.19 | 0.409 | 1.13±0.45 | 1.30±0.60 | 0.562 |
| Glu (mmol/l)a | 5.17±0.20 | 4.95±0.22 | 0.445 | 5.17±0.64 | 4.95±0.76 | 0.565 |
| $E_2$ (nmol/l)b | 0.35±0.07 | 0.16±0.03 | **0.019** | 0.33±0.24 | 0.15±0.11 | *0.085* |
| FSH (IU/l)a | 57.51±11.01 | 94.93±14.14 | 0.053 | 58.31±36.82 | 89.81±39.00 | 0.146 |

BMI=body mass index, LTPA=leisure-time physical activity, HDL-C=high-density lipoprotein cholesterol, LDL-C=low-density lipoprotein cholesterol, TC=total cholesterol, TG=triglycerides, Glu=Fasting blood glucose, $E_2$=estradiol, FSH=follicle stimulating hormone. [a]Statistical analysis run via Student's T-test, [b]Statistical analysis run via Mann-Whitney U-test

**Table S2**: Relative (%) abundancies of sRNA mapping and non-host and host sRNA species in EVs and HDL particles.

| **Time point** | **PRE** | | **POST** | | **1h POST** | |
|---|---|---|---|---|---|---|
| **Carrier particle** | **EV** | **HDL** | **EV** | **HDL** | **EV** | **HDL** |
| ***sRNA mapping*** | | | | | | |
| Mapped to host | 12.0 | 4.5 | | | | |
| Mapped to nonhost | 28.2 | 23.5 | | | | |
| Unmapped | 28.6 | 40.4 | | | | |
| Too Short for Mapping | 31.1 | 31.6 | | | | |
| ***Nonhost sRNA species*** | | | | | | |
| Nonhost rDR | 35.2 | 36.1 | | | | |
| Microbiome bacteria | 12.0 | 13.1 | | | | |
| Algae | 5.6 | 6.0 | | | | |
| Fungus | 3.0 | 3.2 | | | | |
| Nonhost tDR | 2.3 | 2.0 | | | | |
| Environment bacteria | 0.8 | 0.7 | | | | |
| Virus | 0.2 | 0.0 | | | | |
| Mapped to more than one category | 41.0 | 38.8 | | | | |
| ***Host sRNA species*** | | | | | | |
| miR | 51.2 | 29.3 | 54.3 | 28.3 | 46.8 | 21.4 |
| rDR | 34.8 | 49.9 | 29.8 | 37.1 | 33.4 | 48.2 |
| tDR | 12.1 | 18.5 | 14.2 | 32.2 | 17.8 | 27.1 |
| *Other sRNA species* | | | | | | |
| snDR | 0.5 | 0.7 | 0.4 | 0.8 | 0.5 | 0.9 |
| lncDR | 0.4 | 1 | 0.4 | 0.5 | 0.6 | 0.9 |
| mt_tDR | 0.4 | 0.2 | 0.2 | 0.4 | 0.3 | 0.5 |
| yDR | 0.2 | 0.1 | 0.2 | 0.2 | 0.2 | 0.2 |
| snoDR | 0.1 | 0.2 | 0.1 | 0.1 | 0.2 | 0.4 |
| miscDR | 0.3 | 0.3 | 0.3 | 0.4 | 0.4 | 0.5 |

lncDR=long non-coding RNA (lncRNA) –derived sRNA, miR=microRNA, miscRNA=miscellaneous RNA, mt_tRNA= mitochondrial transfer RNA (tRNA) -derived sRNA, rDR=ribosomal RNA (rRNA) -derived sRNA, snoRNA= small nucleolar RNA (snoRNA) –derived sRNA (snoDR), snRNA=small nuclear RNA (snRNA) –derived sRNA (snDR), tDR=transfer RNA (tRNA) -derived sRNA, yDR=Y RNA-derived sRNAs (yDR), miscDR=miscellaneous RNA -derived sRNAs.

**Table S3.** miR counts in EVs and HDL particles in HT users and nonusers at three different time points.

| | | EV | | | | | | HDL | | | | | |
|---|---|---|---|---|---|---|---|---|---|---|---|---|---|
| | | HT user | | | Nonuser | | | HT user | | | Nonuser | | |
| Number | miR | PRE | POST | 1h POST | PRE | POST | 1h POST | PRE | POST | 1h POST | PRE | POST | 1h POST |
| 1 | let-7a-5p | 49402 | 131190 | 40580 | 20209 | 15150 | 8254 | 5831 | 23837 | 6343 | 2307 | 7791 | 4529 |
| 2 | let-7b-5p | 32295 | 78034 | 33114 | 16063 | 11281 | 7708 | 9294 | 11093 | 4093 | 1885 | 5001 | 2156 |
| 3 | let-7c-5p | 1543 | 4498 | 1338 | 855 | 905 | 785 | 71 | 1401 | 359 | 188 | 360 | 100 |
| 4 | let-7d-3p | 80 | 645 | 245 | 71 | 26 | 12 | | | | | | |
| 5 | let-7d-5p | 913 | 5058 | 1023 | 385 | 355 | 172 | 123 | 1191 | 246 | 23 | 107 | 107 |
| 6 | let-7e-5p | 1299 | 5424 | 1729 | 473 | 538 | 230 | 46 | 656 | 166 | 94 | 251 | 92 |
| 7 | let-7f-5p | 24388 | 74443 | 20186 | 11977 | 1063 | 731 | 2750 | 17147 | 6841 | 1592 | 6048 | 4216 |
| 8 | let-7g-5p | 3361 | 13793 | 2629 | 1404 | 3469 | 1968 | 1263 | 1373 | 455 | 141 | 167 | 225 |
| 9 | let-7i-5p | 10259 | 24089 | 8978 | 4906 | 3469 | 1968 | 3646 | 1888 | 812 | 368 | 1066 | 368 |
| 10 | miR-1-3p | 71 | 236 | 118 | 58 | 60 | 109 | 87 | 314 | 152 | 63 | 65 | 97 |
| 11 | miR-7-5p | 141 | 940 | 159 | 225 | 268 | 51 | 172 | 220 | 41 | 16 | 117 | 44 |
| 12 | miR-9-5p | 15 | 160 | 147 | 2 | 32 | 37 | | | | | | |
| 13 | miR-10a-5p | 471 | 7307 | 1441 | 342 | 425 | 192 | 6 | 405 | 100 | 1 | 60 | 127 |
| 14 | miR-10b-5p | 788 | 11072 | 1732 | 314 | 621 | 228 | 2 | 284 | 182 | 7 | 112 | 86 |
| 15 | miR-15a-5p | 213 | 2527 | 302 | 161 | 1003 | 100 | | | | | | |
| 16 | miR-15b-5p | 155 | 1741 | 242 | 226 | 108 | 4 | | | | | | |
| 17 | miR-16-5p | 92140 | 336151 | 113439 | 57850 | 50451 | 16189 | 49867 | 13382 | 7711 | 3735 | 5698 | 4136 |
| 18 | miR-17-5p | 113 | 731 | 190 | 60 | 75 | 54 | | | | | | |
| 19 | miR-18a-5p | 18 | 287 | 4 | 2 | 14 | 0 | | | | | | |
| 20 | miR-19b-3p | 172 | 2878 | 397 | 189 | 146 | 80 | | | | | | |
| 21 | miR-20a-5p | 772 | 6440 | 1113 | 405 | 387 | 135 | 29 | 1574 | 463 | 65 | 119 | 280 |
| 22 | miR-20b-5p | 52 | 824 | 60 | 41 | 182 | 16 | | | | | | |
| 23 | miR-21-5p | 3222 | 23547 | 3675 | 1815 | 2074 | 815 | 1468 | 3192 | 1320 | 430 | 1680 | 1698 |
| 24 | miR-22-3p | 337 | 1780 | 343 | 141 | 72 | 43 | | | | | | |
| 25 | miR-23a-3p | 493 | 4741 | 516 | 205 | 639 | 152 | 625 | 655 | 150 | 44 | 28 | 77 |
| 26 | miR-23b-3p | 1015 | 7711 | 1448 | 643 | 516 | 200 | 79 | 1379 | 339 | 68 | 76 | 200 |
| 27 | miR-24-3p | 1487 | 9170 | 1704 | 547 | 895 | 430 | 552 | 326 | 70 | 23 | 98 | 133 |
| 28 | miR-25-3p | 1008 | 8749 | 2210 | 1107 | 1307 | 304 | 1167 | 803 | 320 | 160 | 256 | 317 |
| 29 | miR-26a-5p | 16665 | 99881 | 18974 | 8919 | 8679 | 3545 | 1830 | 15296 | 4182 | 985 | 3027 | 4103 |

| | | | | | | | | | | | | | |
|---|---|---|---|---|---|---|---|---|---|---|---|---|---|
| 30 | miR-26b-5p | 2034 | 6879 | 1945 | 1230 | 959 | 616 | 278 | 954 | 265 | 96 | 340 | 301 |
| 31 | miR-27a-3p | 211 | 2444 | 282 | 74 | 121 | 63 | | | | | | |
| 32 | miR-27a-5p | 21 | 58 | 67 | 0 | 44 | 92 | | | | | | |
| 33 | miR-27b-3p | 684 | 4319 | 739 | 567 | 299 | 158 | 31 | 119 | 44 | 43 | 277 | 85 |
| 34 | miR-28-3p | 205 | 1888 | 398 | 254 | 700 | 48 | 15 | 461 | 268 | 6 | 89 | 69 |
| 35 | miR-29a-3p | 657 | 5698 | 1091 | 393 | 739 | 113 | 48 | 62 | 8 | 29 | 241 | 82 |
| 36 | miR-29b-3p | 394 | 3743 | 425 | 119 | 407 | 83 | | | | | | |
| 37 | miR-29c-3p | 351 | 4718 | 411 | 648 | 618 | 156 | 149 | 142 | 24 | 43 | 116 | 60 |
| 38 | miR-30a-5p | 1040 | 14018 | 2099 | 682 | 672 | 401 | 302 | 298 | 100 | 17 | 109 | 67 |
| 39 | miR-30b-5p | 61 | 1176 | 70 | 5 | 44 | 45 | | | | | | |
| 40 | miR-30c-5p | 200 | 4039 | 559 | 76 | 94 | 49 | | | | | | |
| 41 | miR-30d-5p | 1691 | 14603 | 3117 | 638 | 898 | 358 | 361 | 1091 | 356 | 84 | 334 | 355 |
| 42 | miR-30e-3p | 86 | 818 | 151 | 29 | 13 | 59 | | | | | | |
| 43 | miR-30e-5p | 1928 | 14138 | 2412 | 1156 | 724 | 377 | 323 | 510 | 143 | 79 | 210 | 185 |
| 44 | miR-32-5p | 34 | 273 | 61 | 8 | 31 | 24 | | | | | | |
| 45 | miR-34a-5p | 41 | 308 | 45 | 55 | 22 | 33 | | | | | | |
| 46 | miR-92a-3p | 3804 | 19689 | 8579 | 1538 | 2237 | 1250 | 1319 | 2851 | 903 | 159 | 1406 | 1236 |
| 47 | miR-92b-3p | 33 | 143 | 0 | 7 | 118 | 52 | | | | | | |
| 48 | miR-92b-5p | 13 | 48 | 8 | 0 | 86 | 52 | | | | | | |
| 49 | miR-93-5p | 2840 | 22624 | 4861 | 3069 | 3430 | 617 | 1032 | 1459 | 1262 | 122 | 683 | 520 |
| 50 | miR-98-5p | 236 | 623 | 169 | 73 | 66 | 45 | 37 | 379 | 39 | 34 | 56 | 50 |
| 51 | miR-99b-5p | 319 | 2760 | 555 | 121 | 90 | 20 | | | | | | |
| 52 | miR-101-3p | 140 | 1753 | 245 | 220 | 157 | 84 | | | | | | |
| 53 | miR-103a-3p | 2453 | 17428 | 4624 | 1292 | 1534 | 416 | 284 | 1864 | 661 | 108 | 459 | 443 |
| 54 | miR-103b | 888 | 6301 | 1698 | 520 | 534 | 145 | 93 | 608 | 219 | 46 | 162 | 128 |
| 55 | miR-107 | 106 | 700 | 70 | 134 | 45 | 38 | | | | | | |
| 56 | miR-122-5p | 8505 | 55628 | 8888 | 16625 | 7537 | 3757 | 20611 | 2375 | 1771 | 1299 | 1120 | 850 |
| 57 | miR-122b-3p | 468 | 2464 | 346 | 1050 | 595 | 244 | 4398 | 163 | 95 | 226 | 60 | 71 |
| 58 | miR-125a-5p | 1569 | 23577 | 5196 | 1272 | 2237 | 492 | 10 | 908 | 477 | 67 | 251 | 240 |
| 59 | miR-125b-5p | 483 | 7690 | 1159 | 798 | 806 | 221 | 19 | 1348 | 246 | 61 | 47 | 94 |
| 60 | miR-126-3p | 52430 | 361333 | 77423 | 56047 | 45292 | 11086 | 4985 | 7491 | 3293 | 465 | 1692 | 1875 |
| 61 | miR-126-5p | 4699 | 48467 | 7165 | 4594 | 3898 | 1155 | 583 | 953 | 863 | 72 | 194 | 276 |
| 62 | miR-127-3p | 40 | 115 | 68 | 3 | 3 | 5 | | | | | | |

| | | | | | | | | | | | | | |
|---|---|---|---|---|---|---|---|---|---|---|---|---|---|
| 63 | miR-128-3p | 101 | 418 | 122 | 19 | 317 | 27 | | | | | | |
| 64 | miR-134-5p | 98 | 196 | 119 | 0 | 12 | 0 | | | | | | |
| 65 | miR-139-3p | 65 | 337 | 172 | 42 | 15 | 16 | | | | | | |
| 66 | miR-139-5p | 69 | 540 | 45 | 26 | 26 | 2 | | | | | | |
| 67 | miR-140-5p | 66 | 227 | 39 | 20 | 26 | 17 | | | | | | |
| 68 | miR-142-3p | 11473 | 58340 | 12103 | 7537 | 9102 | 2759 | 2309 | 9339 | 2243 | 660 | 1633 | 1978 |
| 69 | miR-142-5p | 1172 | 10342 | 1307 | 1622 | 1480 | 246 | | | | | | |
| 70 | miR-143-3p | 380 | 2833 | 577 | 115 | 135 | 80 | 191 | 364 | 121 | 31 | 71 | 80 |
| 71 | miR-144-3p | 151 | 1248 | 334 | 230 | 89 | 47 | | | | | | |
| 72 | miR-144-5p | 254 | 2887 | 522 | 405 | 25 | 27 | | | | | | |
| 73 | miR-146a-5p | 1993 | 12573 | 2460 | 1208 | 1513 | 366 | 566 | 1598 | 862 | 112 | 364 | 367 |
| 74 | miR-146b-5p | 622 | 7197 | 1100 | 682 | 139 | 103 | 45 | 433 | 275 | 60 | 118 | 169 |
| 75 | miR-148a-3p | 625 | 2399 | 605 | 744 | 693 | 185 | 970 | 398 | 350 | 166 | 691 | 228 |
| 76 | miR-148b-3p | 389 | 1946 | 644 | 178 | 486 | 179 | 493 | 833 | 248 | 58 | 144 | 175 |
| 77 | miR-150-5p | 5695 | 63931 | 10089 | 3846 | 5145 | 1685 | 744 | 2424 | 637 | 152 | 354 | 419 |
| 78 | miR-151a-3p | 421 | 1844 | 1097 | 453 | 303 | 72 | 102 | 723 | 523 | 90 | 226 | 116 |
| 79 | miR-155-5p | 856 | 3418 | 1316 | 623 | 496 | 245 | 148 | 107 | 83 | 17 | 39 | 97 |
| 80 | miR-181a-5p | 479 | 3100 | 664 | 143 | 148 | 62 | | | | | | |
| 81 | miR-181b-5p | 69 | 372 | 44 | 46 | 12 | 3 | | | | | | |
| 82 | miR-182-5p | 382 | 1387 | 645 | 115 | 139 | 16 | | | | | | |
| 83 | miR-183-5p | 100 | 509 | 163 | 15 | 163 | 43 | | | | | | |
| 84 | miR-184 | 65 | 93 | 128 | 270 | 49 | 889 | 8 | 799 | 671 | 276 | 148 | 64 |
| 85 | miR-185-5p | 557 | 4305 | 962 | 131 | 215 | 86 | 87 | 1043 | 361 | 35 | 159 | 151 |
| 86 | miR-186-5p | 311 | 2219 | 335 | 138 | 137 | 72 | | | | | | |
| 87 | miR-190a-5p | 194 | 1480 | 206 | 190 | 40 | 44 | | | | | | |
| 88 | miR-191-5p | 1347 | 17649 | 3197 | 3122 | 3792 | 417 | 326 | 3670 | 1383 | 130 | 1059 | 716 |
| 89 | miR-192-5p | 95 | 926 | 74 | 214 | 44 | 45 | | | | | | |
| 90 | miR-194-5p | 61 | 1316 | 107 | 73 | 161 | 150 | | | | | | |
| 91 | miR-196b-5p | 22 | 124 | 10 | 5 | 128 | 39 | | | | | | |
| 92 | miR-197-3p | 76 | 725 | 182 | 5 | 36 | 16 | 22 | 306 | 63 | 2 | 64 | 42 |
| 93 | miR-199a-3p | 650 | 3312 | 933 | 447 | 380 | 127 | 675 | 664 | 392 | 110 | 334 | 215 |
| 94 | miR-199b-3p | 708 | 3129 | 967 | 430 | 324 | 119 | 690 | 740 | 362 | 103 | 344 | 232 |
| 95 | miR-203a-3p | 112 | 160 | 10 | 16 | 16 | 35 | | | | | | |

| | | | | | | | | | | | | | |
|---|---|---|---|---|---|---|---|---|---|---|---|---|---|
| 96 | miR-206 | 7 | 141 | 82 | 149 | 57 | 14 | 7 | 450 | 89 | 229 | 28 | 83 |
| 97 | miR-221-3p | 768 | 4734 | 1190 | 285 | 139 | 77 | 227 | 395 | 105 | 86 | 140 | 217 |
| 98 | miR-222-3p | 66 | 306 | 70 | 25 | 29 | 13 | | | | | | |
| 99 | miR-223-3p | 20442 | 118736 | 25297 | 9664 | 11597 | 3927 | 7826 | 23350 | 5879 | 780 | 2830 | 3547 |
| 100 | miR-223-5p | 172 | 702 | 332 | 256 | 121 | 19 | | | | | | |
| 101 | miR-224-5p | 218 | 515 | 310 | 31 | 7 | 16 | | | | | | |
| 102 | miR-301a-3p | 29 | 298 | 28 | 5 | 19 | 20 | | | | | | |
| 103 | miR-320a-3p | 812 | 3976 | 1518 | 284 | 440 | 196 | 88 | 696 | 94 | 62 | 54 | 131 |
| 104 | miR-320b | 43 | 284 | 64 | 1 | 22 | 43 | | | | | | |
| 105 | miR-328-3p | 104 | 1007 | 675 | 25 | 14 | 18 | 14 | 374 | 88 | 9 | 78 | 98 |
| 106 | miR-335-5p | 216 | 1767 | 407 | 321 | 318 | 93 | 56 | 1531 | 718 | 64 | 150 | 246 |
| 107 | miR-339-5p | 43 | 983 | 385 | 27 | 45 | 17 | | | | | | |
| 108 | miR-340-5p | 165 | 1027 | 106 | 17 | 84 | 17 | | | | | | |
| 109 | miR-342-3p | 2455 | 24690 | 5035 | 2432 | 3479 | 817 | 651 | 1870 | 668 | 24 | 552 | 370 |
| 110 | miR-361-3p | 73 | 1654 | 83 | 46 | 317 | 27 | | | | | | |
| 111 | miR-361-5p | 262 | 2654 | 378 | 86 | 149 | 103 | 11 | 253 | 116 | 5 | 81 | 65 |
| 112 | miR-374a-3p | 23 | 156 | 26 | 0 | 157 | 57 | | | | | | |
| 113 | miR-374a-5p | 228 | 2016 | 401 | 127 | 103 | 32 | | | | | | |
| 114 | miR-379-5p | 36 | 52 | 26 | 19 | 29 | 13 | | | | | | |
| 115 | miR-382-5p | 190 | 365 | 300 | 25 | 46 | 10 | 26 | 127 | 39 | 12 | 38 | 0 |
| 116 | miR-423-5p | 1826 | 4192 | 3097 | 725 | 522 | 397 | 859 | 4966 | 775 | 222 | 1113 | 760 |
| 117 | miR-425-5p | 558 | 3412 | 1864 | 177 | 498 | 200 | 91 | 1063 | 360 | 81 | 398 | 272 |
| 118 | miR-423-3p | 256 | 614 | 346 | 40 | 92 | 25 | | | | | | |
| 119 | miR-429 | | | | | | | 12 | 0 | 2 | 51 | 43 | 20 |
| 120 | miR-432-5p | 221 | 261 | 420 | 33 | 13 | 22 | 55 | 554 | 58 | 19 | 79 | 24 |
| 121 | miR-451a | 2711 | 22341 | 3817 | 1060 | 1503 | 1031 | 541 | 977 | 451 | 56 | 70 | 315 |
| 122 | miR-454-3p | 356 | 1893 | 439 | 129 | 94 | 80 | | | | | | |
| 123 | miR-483-3p | 20 | 273 | 14 | 24 | 43 | 37 | | | | | | |
| 124 | miR-483-5p | 290 | 776 | 509 | 72 | 223 | 89 | | | | | | |
| 125 | miR-486-3p | 45 | 276 | 193 | 16 | 62 | 26 | 4000 | 3848 | 2185 | 471 | 1256 | 1226 |
| 126 | miR-486-5p | 7742 | 35258 | 20108 | 4116 | 5774 | 2749 | | | | | | |
| 127 | miR-574-3p | 4 | 290 | 135 | 13 | 13 | 0 | | | | | | |
| 128 | miR-574-5p | 16 | 64 | 48 | 23 | 10 | 5 | | | | | | |

| | | | | | | | | | | | | | |
|---|---|---|---|---|---|---|---|---|---|---|---|---|---|
| 129 | miR-584-5p | 198 | 703 | 464 | 11 | 83 | 57 | 16 | 188 | 38 | 6 | 35 | 46 |
| 130 | miR-625-3p | 60 | 614 | 363 | 28 | 54 | 31 | | | | | | |
| 131 | miR-654-3p | 15 | 123 | 64 | 0 | 12 | 9 | | | | | | |
| 132 | miR-664a-5p | | | | | | | 3 | 64 | 6 | 0 | 7 | 28 |
| 133 | miR-671-5p | 38 | 99 | 139 | 10 | 2 | 0 | | | | | | |
| 134 | miR-744-5p | 191 | 232 | 455 | 117 | 49 | 53 | 21 | 396 | 27 | 35 | 136 | 47 |
| 135 | miR-769-5p | 39 | 271 | 101 | 33 | 125 | 68 | | | | | | |
| 136 | miR-1277-5p | 7 | 176 | 2 | 8 | 13 | 10 | | | | | | |
| 137 | miR-1307-3p | 104 | 274 | 277 | 34 | 294 | 210 | | | | | | |
| 138 | miR-3065-5p | 34 | 682 | 111 | 125 | 720 | 200 | | | | | | |
| 139 | miR-3135b | 103 | 479 | 285 | 39 | 14 | 26 | 128 | 706 | 372 | 59 | 321 | 601 |
| 140 | miR-3184-3p | 599 | 1420 | 1016 | 256 | 161 | 150 | 302 | 1665 | 247 | 68 | 379 | 263 |
| 141 | miR-3529-3p | 46 | 302 | 65 | 77 | 80 | 18 | 54 | 94 | 23 | 10 | 44 | 16 |
| 142 | miR-3613-5p | 31 | 388 | 13 | 33 | 102 | 20 | | | | | | |
| 143 | miR-4433b-5p | 14 | 79 | 25 | 2 | 58 | 28 | | | | | | |

miR= microRNA

**Table S4**: Twenty most significantly changed miRs in HT users carried via EVs.

| EV | Number | miR | log2 FoldChange | SE (log FoldChange) | Adjusted p-value |
|---|---|---|---|---|---|
| **HT user** | 1 | **miR-191-5p** | 1.190 | 0.285 | **0.004** |
| **POST vs. PRE** | 2 | **miR-223-3p** | 0.681 | 0.172 | **0.005** |
| | 3 | **miR-146b-5p** | 1.143 | 0.336 | **0.032** |
| | 4 | miR-150-5p | 0.523 | 0.173 | 0.091 |
| | 5 | miR-122-5p | -0.667 | 0.231 | 0.109 |
| | 6 | let-7i-5p | -0.555 | 0.208 | 0.151 |
| | 7 | miR-671-5p | -4.109 | 1.522 | 0.151 |
| | 8 | miR-29a-3p | 1.148 | 0.453 | 0.198 |
| | 9 | miR-574-3p | 4.408 | 1.774 | 0.203 |
| | 10 | miR-10a-5p | 0.909 | 0.387 | 0.206 |
| | 11 | miR-125b-5p | 1.158 | 0.479 | 0.206 |
| | 12 | miR-142-3p | 0.346 | 0.147 | 0.206 |
| | 13 | miR-30c-5p | 1.088 | 0.460 | 0.206 |
| | 14 | miR-483-5p | -1.166 | 0.530 | 0.278 |
| | 15 | miR-29c-3p | 1.107 | 0.509 | 0.278 |
| | 16 | miR-486-5p | -0.480 | 0.226 | 0.298 |
| | 17 | miR-27a-5p | -3.414 | 1.697 | 0.362 |
| | 18 | miR-9-5p | 2.710 | 1.360 | 0.362 |
| | 19 | miR-25-3p | 0.617 | 0.322 | 0.408 |
| | 20 | miR-425-5p | 0.759 | 0.418 | 0.485 |
| **HT user** | 1 | **miR-191-5p** | 1.314 | 0.302 | **0.002** |
| **1h POST vs. PRE** | 2 | miR-192-5p | -2.702 | 1.066 | 0.279 |
| | 3 | miR-25-3p | 0.975 | 0.340 | 0.279 |
| | 4 | miR-1-3p | 2.527 | 1.005 | 0.279 |
| | 5 | miR-340-5p | -2.560 | 1.008 | 0.279 |
| | 6 | miR-9-5p | 3.897 | 1.447 | 0.279 |
| | 7 | miR-150-5p | 0.387 | 0.185 | 0.465 |
| | 8 | miR-10a-5p | 0.859 | 0.410 | 0.465 |
| | 9 | miR-574-3p | 4.180 | 1.889 | 0.465 |
| | 10 | miR-92b-3p | -4.367 | 2.005 | 0.465 |
| | 11 | miR-184 | 2.394 | 1.142 | 0.465 |
| | 12 | miR-125b-5p | 1.038 | 0.515 | 0.513 |
| | 13 | miR-223-3p | 0.360 | 0.183 | 0.526 |
| | 14 | miR-103a-3p | 0.432 | 0.226 | 0.526 |
| | 15 | miR-574-5p | 2.232 | 1.153 | 0.526 |
| | 16 | miR-339-5p | 2.324 | 1.235 | 0.528 |
| | 17 | let-7f-5p | -0.290 | 0.171 | 0.581 |
| | 18 | let-7g-5p | -0.402 | 0.271 | 0.581 |
| | 19 | miR-30d-5p | -0.485 | 0.348 | 0.581 |
| | 20 | miR-425-5p | 0.720 | 0.447 | 0.581 |
| **HT user** | 1 | **miR-486-5p** | 0.869 | 0.240 | **0.041** |
| **1h POST vs. POST** | 2 | **miR-340-5p** | -3.432 | 0.998 | **0.041** |
| | 3 | miR-146b-5p | -0.816 | 0.352 | 0.419 |
| | 4 | miR-744-5p | 2.092 | 0.854 | 0.419 |
| | 5 | miR-1-3p | 2.294 | 0.992 | 0.419 |
| | 6 | miR-184 | 2.789 | 1.140 | 0.419 |
| | 7 | miR-671-5p | 3.869 | 1.619 | 0.419 |
| | 8 | miR-103a-3p | 0.491 | 0.225 | 0.517 |
| | 9 | miR-29a-3p | -1.008 | 0.476 | 0.522 |
| | 10 | miR-374a-5p | 1.828 | 0.877 | 0.522 |
| | 11 | miR-122-5p | 0.396 | 0.246 | 0.595 |

| | | | | |
|---|---|---|---|---|
| 12 | miR-223-3p | -0.322 | 0.182 | 0.595 |
| 13 | miR-26a-5p | -0.327 | 0.179 | 0.595 |
| 14 | miR-3184-3p | 0.553 | 0.374 | 0.595 |
| 15 | miR-423-5p | 0.460 | 0.311 | 0.595 |
| 16 | miR-483-5p | 1.082 | 0.560 | 0.595 |
| 17 | miR-92a-3p | 0.378 | 0.224 | 0.595 |
| 18 | miR-103b | 0.512 | 0.296 | 0.595 |
| 19 | miR-10b-5p | -0.941 | 0.544 | 0.595 |
| 20 | miR-146a-5p | -0.435 | 0.277 | 0.595 |

miR=microRNA, HT=hormone therapy. Statistically significant findings are marked in bold.

**Table S5**: Twenty most significantly changed miRs in nonusers carried via EVs.

| EV | Number | miR | log2 FoldChange | SE (log FoldChange) | Adjusted p-value |
|---|---|---|---|---|---|
| **Nonuser** | 1 | miR-122-5p | -1.259 | 0.440 | 0.594 |
| **POST vs. PRE** | 2 | miR-186-5p | -2.822 | 1.159 | 0.702 |
| | 3 | miR-190a-5p | -4.039 | 1.628 | 0.702 |
| | 4 | miR-122b-3p | -1.362 | 0.614 | 0.829 |
| | 5 | miR-192-5p | -2.983 | 1.370 | 0.829 |
| | 6 | let-7a-5p | -0.052 | 0.258 | 0.997 |
| | 7 | let-7b-5p | -0.290 | 0.275 | 0.997 |
| | 8 | let-7c-5p | 0.306 | 0.692 | 0.997 |
| | 9 | let-7d-5p | -0.573 | 0.702 | 0.997 |
| | 10 | let-7f-5p | -0.045 | 0.227 | 0.997 |
| | 11 | let-7g-5p | -0.851 | 0.503 | 0.997 |
| | 12 | let-7i-5p | -0.174 | 0.323 | 0.997 |
| | 13 | miR-126-3p | 0.132 | 0.319 | 0.997 |
| | 14 | miR-146b-5p | 0.347 | 0.889 | 0.997 |
| | 15 | miR-148a-3p | -0.824 | 0.747 | 0.997 |
| | 16 | miR-150-5p | 0.001 | 0.281 | 0.997 |
| | 17 | miR-16-5p | -0.051 | 0.237 | 0.997 |
| | 18 | miR-17-5p | 1.132 | 1.565 | 0.997 |
| | 19 | miR-191-5p | 0.873 | 0.498 | 0.997 |
| | 20 | miR-199a-3p | -0.295 | 0.621 | 0.997 |
| **Nonuser** | 1 | let-7a-5p | 0.086 | 0.261 | 0.921 |
| **1h POST vs. PRE** | 2 | let-7b-5p | 0.205 | 0.277 | 0.921 |
| | 3 | let-7c-5p | 0.369 | 0.703 | 0.921 |
| | 4 | let-7d-5p | -0.608 | 0.716 | 0.921 |
| | 5 | let-7f-5p | -0.279 | 0.231 | 0.921 |
| | 6 | let-7g-5p | -0.384 | 0.524 | 0.921 |
| | 7 | let-7i-5p | 0.130 | 0.326 | 0.921 |
| | 8 | miR-122-5p | -0.688 | 0.441 | 0.921 |
| | 9 | miR-126-3p | -0.271 | 0.319 | 0.921 |
| | 10 | miR-146b-5p | -0.885 | 0.917 | 0.921 |
| | 11 | miR-148a-3p | -0.783 | 0.765 | 0.921 |
| | 12 | miR-150-5p | -0.200 | 0.286 | 0.921 |
| | 13 | miR-16-5p | -0.230 | 0.237 | 0.921 |
| | 14 | miR-17-5p | 2.276 | 1.568 | 0.921 |
| | 15 | miR-186-5p | -2.364 | 1.189 | 0.921 |
| | 16 | miR-190a-5p | -2.228 | 1.605 | 0.921 |
| | 17 | miR-191-5p | -0.328 | 0.508 | 0.921 |
| | 18 | miR-199b-3p | -0.215 | 0.628 | 0.921 |
| | 19 | miR-223-3p | 0.265 | 0.316 | 0.921 |
| | 20 | miR-223-5p | -2.186 | 1.429 | 0.921 |
| **Nonuser** | 1 | miR-191-5p | -1.201 | 0.502 | 0.854 |
| **1h POST vs. POST** | 2 | miR-28-3p | -2.300 | 1.001 | 0.854 |
| | 3 | miR-107 | 3.114 | 1.488 | 0.854 |
| | 4 | miR-142-3p | -0.470 | 0.211 | 0.854 |
| | 5 | miR-432-5p | 4.287 | 2.042 | 0.854 |
| | 6 | miR-181a-5p | 3.349 | 1.339 | 0.854 |
| | 7 | let-7b-5p | 0.495 | 0.276 | 0.928 |
| | 8 | miR-342-3p | -0.805 | 0.419 | 0.928 |
| | 9 | miR-451a | 1.205 | 0.627 | 0.928 |
| | 10 | miR-361-3p | -2.872 | 1.532 | 0.928 |

| | | | | |
|---|---|---|---|---|
| 11 | miR-184 | 3.101 | 1.706 | 0.928 |
| 12 | let-7a-5p | 0.138 | 0.260 | 0.991 |
| 13 | let-7c-5p | 0.062 | 0.693 | 0.991 |
| 14 | let-7d-5p | -0.035 | 0.713 | 0.991 |
| 15 | let-7f-5p | -0.234 | 0.229 | 0.991 |
| 16 | let-7g-5p | 0.467 | 0.521 | 0.991 |
| 17 | let-7i-5p | 0.303 | 0.322 | 0.991 |
| 18 | miR-122-5p | 0.570 | 0.441 | 0.991 |
| 19 | miR-126-3p | -0.402 | 0.319 | 0.991 |
| 20 | miR-146b-5p | -1.232 | 0.911 | 0.991 |

miR=microRNA.

**Table S6**: Twenty most significantly changed miRs in HT users carried via HDL.

| HDL | Number | miR | log2 FoldChange | SE (log FoldChange) | Adjusted p-value |
|---|---|---|---|---|---|
| **HT user** | 1 | **let-7c-5p** | 3.450 | 0.680 | **0.000** |
| **POST vs. PRE** | 2 | **miR-125b-5p** | 5.195 | 1.294 | **0.001** |
| | 3 | **miR-206** | 4.988 | 1.215 | **0.001** |
| | 4 | **miR-184** | 6.466 | 1.758 | **0.004** |
| | 5 | miR-26a-5p | 1.161 | 0.431 | 0.106 |
| | 6 | miR-7-5p | 4.114 | 1.582 | 0.116 |
| | 7 | miR-10a-5p | 4.481 | 1.782 | 0.128 |
| | 8 | miR-125a-5p | 2.929 | 1.249 | 0.159 |
| | 9 | miR-20a-5p | 3.214 | 1.355 | 0.159 |
| | 10 | let-7f-5p | 0.907 | 0.414 | 0.213 |
| | 11 | miR-122b-3p | -2.353 | 1.105 | 0.222 |
| | 12 | miR-3529-3p | 4.192 | 1.994 | 0.222 |
| | 13 | let-7e-5p | 2.051 | 1.066 | 0.313 |
| | 14 | miR-103a-3p | 1.629 | 0.861 | 0.313 |
| | 15 | miR-122-5p | -1.045 | 0.605 | 0.421 |
| | 16 | miR-143-3p | 2.901 | 1.771 | 0.476 |
| | 17 | let-7a-5p | 0.603 | 0.379 | 0.493 |
| | 18 | miR-584-5p | 2.861 | 1.861 | 0.517 |
| | 19 | miR-93-5p | 1.202 | 0.869 | 0.599 |
| | 20 | miR-103b | 1.520 | 1.128 | 0.599 |
| **HT user** | 1 | **miR-206** | 4.903 | 1.216 | **0.003** |
| **1h POST vs. PRE** | 2 | **miR-184** | 6.953 | 1.757 | **0.003** |
| | 3 | **miR-125b-5p** | 4.533 | 1.295 | **0.012** |
| | 4 | **let-7c-5p** | 2.158 | 0.684 | **0.030** |
| | 5 | miR-125a-5p | 3.548 | 1.245 | 0.066 |
| | 6 | miR-93-5p | 2.270 | 0.864 | 0.107 |
| | 7 | miR-20a-5p | 2.893 | 1.360 | 0.359 |
| | 8 | miR-26a-5p | 0.849 | 0.431 | 0.460 |
| | 9 | miR-486-5p | 1.116 | 0.628 | 0.629 |
| | 10 | let-7f-5p | 0.584 | 0.414 | 0.732 |
| | 11 | miR-150-5p | 1.142 | 0.824 | 0.732 |
| | 12 | miR-28-3p | 2.985 | 1.914 | 0.732 |
| | 13 | miR-320a-3p | -2.177 | 1.502 | 0.732 |
| | 14 | miR-451a | 1.347 | 0.855 | 0.732 |
| | 15 | miR-126-5p | 1.203 | 0.860 | 0.732 |
| | 16 | miR-7-5p | 2.335 | 1.593 | 0.732 |
| | 17 | miR-328-3p | 2.865 | 1.853 | 0.732 |
| | 18 | miR-146b-5p | 2.256 | 1.859 | 0.844 |
| | 19 | miR-30a-5p | 1.900 | 1.686 | 0.844 |
| | 20 | miR-103a-3p | 0.969 | 0.868 | 0.844 |
| **HT user** | 1 | miR-486-5p | 1.524 | 0.596 | 0.797 |
| **1h POST vs. POST** | 2 | let-7a-5p | -0.479 | 0.358 | 0.881 |
| | 3 | let-7b-5p | -0.408 | 0.353 | 0.881 |
| | 4 | let-7c-5p | -1.292 | 0.624 | 0.881 |
| | 5 | let-7i-5p | -1.082 | 0.669 | 0.881 |
| | 6 | miR-122-5p | 0.665 | 0.572 | 0.881 |
| | 7 | miR-24-3p | -1.643 | 1.619 | 0.881 |
| | 8 | miR-30a-5p | 2.997 | 1.614 | 0.881 |
| | 9 | miR-30e-5p | 2.814 | 1.556 | 0.881 |
| | 10 | miR-3184-3p | -0.797 | 0.697 | 0.881 |
| | 11 | miR-320a-3p | -2.001 | 1.416 | 0.881 |

| | | | | |
|---|---|---|---|---|
| 12 | miR-423-5p | -0.588 | 0.568 | 0.881 |
| 13 | miR-451a | 1.252 | 0.802 | 0.881 |
| 14 | miR-93-5p | 1.067 | 0.793 | 0.881 |
| 15 | let-7e-5p | -1.329 | 0.985 | 0.881 |
| 16 | miR-10a-5p | -2.308 | 1.605 | 0.881 |
| 17 | miR-122b-3p | 1.269 | 1.052 | 0.881 |
| 18 | miR-146a-5p | 0.840 | 0.684 | 0.881 |
| 19 | miR-23a-3p | 1.374 | 1.374 | 0.881 |
| 20 | miR-25-3p | -1.373 | 1.315 | 0.881 |

miR=microRNA, HT= hormone therapy. Statistically significant findings are marked in bold.

**Table S7**: Twenty most significantly changed miRs in nonusers carried via HDL.

| HDL | Number | miR | log2 FoldChange | SE (log FoldChange) | Adjusted p-value |
|---|---|---|---|---|---|
| **Nonuser** | 1 | miR-28-3p | 6.645 | 2.442 | 0.244 |
| **POST vs. PRE** | 2 | miR-429 | -5.920 | 2.020 | 0.244 |
| | 3 | let-7g-5p | -2.336 | 0.913 | 0.247 |
| | 4 | miR-342-3p | 2.553 | 1.064 | 0.247 |
| | 5 | miR-92a-3p | 1.231 | 0.507 | 0.247 |
| | 6 | let-7d-5p | 1.542 | 1.681 | 0.879 |
| | 7 | let-7f-5p | 0.567 | 0.367 | 0.879 |
| | 8 | let-7i-5p | 0.480 | 0.534 | 0.879 |
| | 9 | miR-122-5p | -0.673 | 0.655 | 0.879 |
| | 10 | miR-146b-5p | 1.398 | 1.261 | 0.879 |
| | 11 | miR-191-5p | 0.719 | 0.641 | 0.879 |
| | 12 | miR-199a-3p | 1.329 | 0.963 | 0.879 |
| | 13 | miR-199b-3p | 0.985 | 0.924 | 0.879 |
| | 14 | miR-23b-3p | -1.441 | 1.358 | 0.879 |
| | 15 | miR-30a-5p | 2.603 | 2.001 | 0.879 |
| | 16 | miR-30e-5p | -1.454 | 0.951 | 0.879 |
| | 17 | miR-320a-3p | -1.999 | 1.172 | 0.879 |
| | 18 | miR-361-5p | 2.085 | 2.011 | 0.879 |
| | 19 | miR-423-5p | 0.502 | 0.595 | 0.879 |
| | 20 | miR-486-5p | -0.538 | 0.489 | 0.879 |
| **Nonuser** | 1 | miR-28-3p | 7.479 | 2.602 | 0.152 |
| **1h POST vs. PRE** | 2 | miR-429 | -6.355 | 2.168 | 0.152 |
| | 3 | miR-342-3p | 2.957 | 1.131 | 0.224 |
| | 4 | miR-92a-3p | 1.268 | 0.540 | 0.353 |
| | 5 | miR-155-5p | 4.220 | 2.130 | 0.713 |
| | 6 | let-7c-5p | -1.827 | 0.999 | 0.716 |
| | 7 | miR-20a-5p | 2.511 | 1.352 | 0.716 |
| | 8 | miR-3135b | 2.465 | 1.391 | 0.716 |
| | 9 | let-7a-5p | -0.214 | 0.394 | 0.966 |
| | 10 | let-7b-5p | -0.300 | 0.331 | 0.966 |
| | 11 | let-7d-5p | 1.812 | 1.807 | 0.966 |
| | 12 | let-7f-5p | 0.415 | 0.393 | 0.966 |
| | 13 | let-7g-5p | -0.157 | 0.966 | 0.966 |
| | 14 | let-7i-5p | -0.342 | 0.578 | 0.966 |
| | 15 | miR-122-5p | -0.413 | 0.699 | 0.966 |
| | 16 | miR-126-3p | 0.750 | 0.577 | 0.966 |
| | 17 | miR-146b-5p | 0.537 | 1.351 | 0.966 |
| | 18 | miR-148a-3p | -1.055 | 0.684 | 0.966 |
| | 19 | miR-150-5p | 0.479 | 1.092 | 0.966 |
| | 20 | miR-16-5p | -0.041 | 0.475 | 0.966 |
| **Nonuser** | 1 | let-7c-5p | -2.082 | 0.979 | 0.815 |
| **1h POST vs. POST** | 2 | let-7g-5p | 2.180 | 0.979 | 0.815 |
| | 3 | miR-148b-3p | 2.228 | 1.130 | 0.815 |
| | 4 | miR-155-5p | 4.839 | 2.119 | 0.815 |
| | 5 | miR-184 | 3.058 | 1.589 | 0.815 |
| | 6 | miR-20a-5p | 2.326 | 1.303 | 0.927 |
| | 7 | let-7a-5p | -0.235 | 0.392 | 0.931 |
| | 8 | let-7b-5p | -0.246 | 0.327 | 0.931 |
| | 9 | let-7d-5p | 0.269 | 1.745 | 0.931 |
| | 10 | let-7f-5p | -0.152 | 0.389 | 0.931 |
| | 11 | let-7i-5p | -0.822 | 0.562 | 0.931 |

| | | | | |
|---|---|---|---|---|
| 12 | miR-122-5p | 0.260 | 0.694 | 0.931 |
| 13 | miR-126-3p | 0.420 | 0.567 | 0.931 |
| 14 | miR-146b-5p | -0.861 | 1.313 | 0.931 |
| 15 | miR-148a-3p | -0.792 | 0.676 | 0.931 |
| 16 | miR-150-5p | 0.780 | 1.077 | 0.931 |
| 17 | miR-16-5p | 0.052 | 0.474 | 0.931 |
| 18 | miR-191-5p | -0.571 | 0.679 | 0.931 |
| 19 | miR-199a-3p | -0.793 | 1.004 | 0.931 |
| 20 | miR-199b-3p | -0.752 | 0.972 | 0.931 |

miR=microRNA.

**Table S8:** Twenty most significantly changed rDR species in HT users carried via EVs.

| EV | Number | rDR | Annotation | log2 FoldChange | SE (log FoldChange) | Adjusted p-value |
|---|---|---|---|---|---|---|
| **HT user** | 1 | URS0001AF915F_9606 | Homo sapiens (bacterial) SSU rRNA | -1.115 | 0.354 | 0.791 |
| **POST vs. PRE** | 2 | URS0001B492FB_9606 | Homo sapiens SSU rRNA | 1.483 | 0.467 | 0.791 |
| | 3 | URS0001B8AA51_9606 | Homo sapiens SSU rRNA | 1.198 | 0.365 | 0.791 |
| | 4 | URS0001BB1A02_9606 | Homo sapiens SSU rRNA | 1.694 | 0.550 | 0.791 |
| | 5 | URS0001AC3CEC_9606 | Homo sapiens SSU rRNA | 1.340 | 0.391 | 0.791 |
| | 6 | URS0000005270_9606 | Homo sapiens RNA, 5.8S ribosomal N3 | -0.141 | 0.300 | 0.996 |
| | 7 | URS000002B0D5_9606 | Homo sapiens 5S rRNA | -0.380 | 0.349 | 0.996 |
| | 8 | URS000005D9C4_9606 | Homo sapiens 28S rRNA V8 region | -0.534 | 0.397 | 0.996 |
| | 9 | URS0000086682_9606 | Homo sapiens 28S rRNA V8 region | -0.486 | 0.382 | 0.996 |
| | 10 | URS00000D0A5B_9606 | Homo sapiens partial rRNA | 0.175 | 0.318 | 0.996 |
| | 11 | URS00000F9D45_9606 | Homo sapiens RNA, 5S ribosomal 12 | -0.388 | 0.344 | 0.996 |
| | 12 | URS00001152CA_9606 | Homo sapiens partial rRNA | 0.273 | 0.295 | 0.996 |
| | 13 | URS000013AA81_9606 | Homo sapiens 28S rRNA V8 region | -0.422 | 0.364 | 0.996 |
| | 14 | URS0000165891_9606 | Homo sapiens SSU rRNA (18S) | -0.260 | 0.252 | 0.996 |
| | 15 | URS000018CAFF_9606 | Homo sapiens 28S rRNA V8 region | -0.521 | 0.365 | 0.996 |
| | 16 | URS000018FE5D_9606 | Homo sapiens LSU rRNA (28S) | 0.151 | 0.284 | 0.996 |
| | 17 | URS000019B9A7_9606 | Homo sapiens 28S rRNA V8 region | -0.154 | 0.379 | 0.996 |
| | 18 | URS00001AFAED_9606 | Homo sapiens partial rRNA | 0.478 | 0.312 | 0.996 |
| | 19 | URS00001C0ED1_9606 | Homo sapiens rRNA SSU 18S | 0.022 | 0.259 | 0.996 |
| | 20 | URS00001E876A_9606 | Homo sapiens 28S rRNA V8 region | -0.359 | 0.339 | 0.996 |
| **HT user** | 1 | **URS0001BB76DE_9606** | Homo sapiens (mitochondrial) SSU rRNA | 1.976 | 0.396 | **0.001** |
| **1h POST vs. PRE** | 2 | **URS0001B96E0D_9606** | Homo sapiens (mitochondrial) SSU rRNA | 1.952 | 0.431 | **0.003** |
| | 3 | **URS0001A68F15_9606** | Homo sapiens (mitochondrial) SSU rRNA | 1.887 | 0.445 | **0.007** |
| | 4 | **URS0001B4DC43_9606** | Homo sapiens (mitochondrial) SSU rRNA | 2.061 | 0.499 | **0.007** |
| | 5 | **URS0001B5B3F4_9606** | Homo sapiens (mitochondrial) SSU rRNA | 1.606 | 0.395 | **0.007** |
| | 6 | **URS0001BAA09A_9606** | Homo sapiens (mitochondrial) SSU rRNA | 2.027 | 0.500 | **0.007** |
| | 7 | **URS00021358C2_9606** | Homo sapiens (mitochondrial) LSU rRNA | 2.476 | 0.623 | **0.009** |
| | 8 | **URS0001A7C0B1_9606** | Homo sapiens (mitochondrial) SSU rRNA | 1.792 | 0.469 | **0.010** |
| | 9 | **URS0001A93ED7_9606** | Homo sapiens (mitochondrial) SSU rRNA | 2.125 | 0.558 | **0.010** |
| | 10 | **URS0001AF034E_9606** | Homo sapiens (mitochondrial) SSU rRNA | 2.609 | 0.685 | **0.010** |
| | 11 | **URS0001B52AC1_9606** | Homo sapiens (mitochondrial) SSU rRNA | 1.546 | 0.404 | **0.010** |
| | 12 | **URS0001A7919B_9606** | Homo sapiens (mitochondrial) SSU rRNA | 1.743 | 0.458 | **0.010** |
| | 13 | **URS0001B175EC_9606** | Homo sapiens (mitochondrial) SSU rRNA | 2.125 | 0.564 | **0.011** |
| | 14 | **URS0001A4865C_9606** | Homo sapiens (mitochondrial) SSU rRNA | 1.748 | 0.473 | **0.014** |

| | | | | | | |
|---|---|---|---|---|---|---|
| | 15 | **URS0001AB4292_9606** | Homo sapiens (mitochondrial) SSU rRNA | 1.928 | 0.530 | **0.016** |
| | 16 | **URS0001AE0470_9606** | Homo sapiens (mitochondrial) SSU rRNA | 1.745 | 0.485 | **0.017** |
| | 17 | **URS0001B81ABB_9606** | Homo sapiens (mitochondrial) SSU rRNA | 1.295 | 0.367 | **0.022** |
| | 18 | **URS0001A5136D_9606** | Homo sapiens (mitochondrial) SSU rRNA | 2.116 | 0.606 | **0.023** |
| | 19 | **URS00021233ED_9606** | Homo sapiens (mitochondrial) LSU rRNA | 1.262 | 0.363 | **0.023** |
| | 20 | **URS0001A2D729_9606** | Homo sapiens (mitochondrial) SSU rRNA | 1.280 | 0.376 | **0.027** |
| **HT user** | 1 | **URS0001A7C0B1_9606** | Homo sapiens (mitochondrial) SSU rRNA | 2.061 | 0.475 | **0.004** |
| **1h POST vs. POST** | 2 | **URS0001B4DC43_9606** | Homo sapiens (mitochondrial) SSU rRNA | 2.188 | 0.508 | **0.004** |
| | 3 | **URS0001BB76DE_9606** | Homo sapiens (mitochondrial) SSU rRNA | 1.655 | 0.382 | **0.004** |
| | 4 | **URS0001BAA09A_9606** | Homo sapiens (mitochondrial) SSU rRNA | 2.243 | 0.497 | **0.004** |
| | 5 | **URS0001B82586_9606** | Homo sapiens (mitochondrial) SSU rRNA | 1.415 | 0.339 | **0.005** |
| | 6 | **URS0001A93ED7_9606** | Homo sapiens (mitochondrial) SSU rRNA | 2.257 | 0.550 | **0.006** |
| | 7 | **URS0001B52AC1_9606** | Homo sapiens (mitochondrial) SSU rRNA | 1.605 | 0.396 | **0.006** |
| | 8 | **URS0001B96E0D_9606** | Homo sapiens (mitochondrial) SSU rRNA | 1.746 | 0.427 | **0.006** |
| | 9 | **URS0001A820F5_9606** | Homo sapiens (mitochondrial) SSU rRNA | 1.840 | 0.457 | **0.006** |
| | 10 | **URS0001AEA987_9606** | Homo sapiens (mitochondrial) SSU rRNA | 1.910 | 0.485 | **0.006** |
| | 11 | **URS0001AF034E_9606** | Homo sapiens (mitochondrial) SSU rRNA | 2.757 | 0.694 | **0.006** |
| | 12 | **URS00021358C2_9606** | Homo sapiens (mitochondrial) LSU rRNA | 2.466 | 0.624 | **0.006** |
| | 13 | **URS0001A4865C_9606** | Homo sapiens (mitochondrial) SSU rRNA | 1.830 | 0.471 | **0.006** |
| | 14 | **URS0001AB447F_9606** | Homo sapiens (mitochondrial) SSU rRNA | 1.106 | 0.289 | **0.006** |
| | 15 | **URS0001AE134F_9606** | Homo sapiens (mitochondrial) SSU rRNA | 1.627 | 0.427 | **0.006** |
| | 16 | **URS0001B5BD37_9606** | Homo sapiens (mitochondrial) SSU rRNA | 2.443 | 0.638 | **0.006** |
| | 17 | **URS0001B959E6_9606** | Homo sapiens (mitochondrial) SSU rRNA | 2.456 | 0.636 | **0.006** |
| | 18 | **URS0001BA4530_9606** | Homo sapiens (mitochondrial) SSU rRNA | 1.399 | 0.363 | **0.006** |
| | 19 | **URS0001A99078_9606** | Homo sapiens (mitochondrial) SSU rRNA | 1.978 | 0.507 | **0.006** |
| | 20 | **URS0001B87590_9606** | Homo sapiens (mitochondrial) SSU rRNA | 2.545 | 0.666 | **0.006** |

rDR=ribosomal RNA (rRNA) -derived sRNA. SSU=small subunit, LSU=large subunit. Statistically significant findings are marked in bold.

**Table S9:** 20 most significantly changed rDR species in nonusers carried via EVs.

| EV | Number | rDR | Annotation | SE (log FoldChange) | Adjusted p-value |
|---|---|---|---|---|---|
| **Nonuser POST vs. PRE** | 1 | **URS0001BA83D1_9606** | Homo sapiens (mitochondrial) SSU rRNA | 0.607 | **0.004** |
| | 2 | **URS0001A53ABA_9606** | Homo sapiens (mitochondrial) SSU rRNA | 0.521 | **0.007** |
| | 3 | **URS0001AF034E_9606** | Homo sapiens (mitochondrial) SSU rRNA | 0.663 | **0.007** |
| | 4 | **URS0001B3F905_9606** | Homo sapiens (mitochondrial) SSU rRNA | 0.659 | **0.007** |
| | 5 | **URS0001BA11BB_9606** | Homo sapiens (mitochondrial) SSU rRNA | 0.629 | **0.007** |
| | 6 | **URS0001B12279_9606** | Homo sapiens (mitochondrial) SSU rRNA | 0.675 | **0.009** |
| | 7 | **URS00021358C2_9606** | Homo sapiens (mitochondrial) LSU rRNA | 0.745 | **0.010** |
| | 8 | **URS0001B38803_9606** | Homo sapiens (mitochondrial) SSU rRNA | 0.627 | **0.011** |
| | 9 | **URS0001B6D42D_9606** | Homo sapiens (mitochondrial) SSU rRNA | 0.560 | **0.011** |
| | 10 | **URS0001B959E6_9606** | Homo sapiens (mitochondrial) SSU rRNA | 0.660 | **0.011** |
| | 11 | **URS0001B5BD37_9606** | Homo sapiens (mitochondrial) SSU rRNA | 0.663 | **0.011** |
| | 12 | **URS0001A3E299_9606** | Homo sapiens (mitochondrial) SSU rRNA | 0.619 | **0.011** |
| | 13 | **URS0001B58777_9606** | Homo sapiens (mitochondrial) SSU rRNA | 0.521 | **0.012** |
| | 14 | **URS0001BBDC10_9606** | Homo sapiens (mitochondrial) SSU rRNA | 0.513 | **0.013** |
| | 15 | **URS0001A293EB_9606** | Homo sapiens (mitochondrial) SSU rRNA | 0.449 | **0.019** |
| | 16 | **URS0001B20D6D_9606** | Homo sapiens (mitochondrial) SSU rRNA | 0.695 | **0.019** |
| | 17 | **URS0001B1FE0C_9606** | Homo sapiens (mitochondrial) SSU rRNA | 0.446 | **0.021** |
| | 18 | **URS0001A54C0A_9606** | Homo sapiens (mitochondrial) SSU rRNA | 0.576 | **0.024** |
| | 19 | **URS0001A93ED7_9606** | Homo sapiens (mitochondrial) SSU rRNA | 0.846 | **0.024** |
| | 20 | **URS0001B0FE9E_9606** | Homo sapiens (mitochondrial) SSU rRNA | 0.641 | **0.024** |
| **Nonuser 1h POST vs. PRE** | 1 | URS00006B81C9_9606 | Homo sapiens 5.8S rRNA | 1.085 | 0.930 |
| | 2 | URS000068483A_9606 | Homo sapiens 5S rRNA | 0.744 | 0.963 |
| | 3 | URS0001A4C3CA_9606 | Homo sapiens SSU rRNA | 0.427 | 0.963 |
| | 4 | URS0000005270_9606 | Homo sapiens RNA, 5.8S ribosomal N3 | 0.424 | 0.984 |
| | 5 | URS00000D0A5B_9606 | Homo sapiens partial rRNA | 0.367 | 0.984 |
| | 6 | URS00001152CA_9606 | Homo sapiens partial rRNA | 0.421 | 0.984 |
| | 7 | URS0000162C88_9606 | Homo sapiens rRNA LSU-S 5.8S | 0.393 | 0.984 |
| | 8 | URS0000171707_9606 | Homo sapiens partial rRNA | 0.418 | 0.984 |
| | 9 | URS000018CAFF_9606 | Homo sapiens 28S rRNA V8 region | 0.436 | 0.984 |
| | 10 | URS000019B9A7_9606 | Homo sapiens 28S rRNA V8 region | 0.507 | 0.984 |
| | 11 | URS00001AFAED_9606 | Homo sapiens partial rRNA | 0.418 | 0.984 |
| | 12 | URS00001C0ED1_9606 | Homo sapiens rRNA SSU 18S | 0.314 | 0.984 |
| | 13 | URS000021E2D0_9606 | Homo sapiens partial rRNA | 0.476 | 0.984 |

| | | | | | |
|---|---|---|---|---|---|
| | 14 | URS000023CAC4_9606 | Homo sapiens SSU rRNA (18S) | 0.642 | 0.984 |
| | 15 | URS000027500B_9606 | Homo sapiens partial rRNA | 0.437 | 0.984 |
| | 16 | URS00002A452D_9606 | Homo sapiens 28S rRNA V8 region | 0.436 | 0.984 |
| | 17 | URS000031689A_9606 | Homo sapiens SSU rRNA (18S) | 0.350 | 0.984 |
| | 18 | URS000032E2CD_9606 | Homo sapiens SSU rRNA (18S) | 0.299 | 0.984 |
| | 19 | URS000034CFF8_9606 | Homo sapiens partial rRNA | 0.480 | 0.984 |
| | 20 | URS0000385E81_9606 | Homo sapiens 28S rRNA V8 region | 0.455 | 0.984 |
| **Nonuser** | 1 | URS0001ABA328_9606 | Homo sapiens (mitochondrial) SSU rRNA | 0.527 | 0.614 |
| **1h POST vs. POST** | 2 | URS00004931F8_9606 | Homo sapiens 28S rRNA V8 region | 0.442 | 0.756 |
| | 3 | URS000052EE02_9606 | Homo sapiens 28S rRNA V8 region | 0.450 | 0.756 |
| | 4 | URS000053F3EB_9606 | Homo sapiens 28S rRNA V8 region | 0.436 | 0.756 |
| | 5 | URS00006E97F5_9606 | Homo sapiens SSU rRNA | 0.474 | 0.756 |
| | 6 | URS00019D4CA7_9606 | Homo sapiens LHRI_LNC27.2 | 0.350 | 0.756 |
| | 7 | URS0001A293EB_9606 | Homo sapiens (mitochondrial) SSU rRNA | 0.441 | 0.756 |
| | 8 | URS0001A3FAC1_9606 | Homo sapiens (mitochondrial) SSU rRNA | 0.571 | 0.756 |
| | 9 | URS0001A53ABA_9606 | Homo sapiens (mitochondrial) SSU rRNA | 0.518 | 0.756 |
| | 10 | URS0001A54C0A_9606 | Homo sapiens (mitochondrial) SSU rRNA | 0.572 | 0.756 |
| | 11 | URS0001A60BBC_9606 | Homo sapiens SSU rRNA | 0.563 | 0.756 |
| | 12 | URS0001AD239E_9606 | Homo sapiens (mitochondrial) SSU rRNA | 0.442 | 0.756 |
| | 13 | URS0001ADA731_9606 | Homo sapiens (mitochondrial) SSU rRNA | 0.459 | 0.756 |
| | 14 | URS0001ADE278_9606 | Homo sapiens (mitochondrial) SSU rRNA | 0.512 | 0.756 |
| | 15 | URS0001AEB7F8_9606 | Homo sapiens SSU rRNA | 0.681 | 0.756 |
| | 16 | URS0001AF034E_9606 | Homo sapiens (mitochondrial) SSU rRNA | 0.658 | 0.756 |
| | 17 | URS0001B00FC7_9606 | Homo sapiens (mitochondrial) SSU rRNA | 0.555 | 0.756 |
| | 18 | URS0001B05F21_9606 | Homo sapiens (mitochondrial) SSU rRNA | 0.461 | 0.756 |
| | 19 | URS0001B1E32A_9606 | Homo sapiens SSU rRNA | 0.303 | 0.756 |
| | 20 | URS0001B20D6D_9606 | Homo sapiens (mitochondrial) SSU rRNA | 0.681 | 0.756 |

rDR= ribosomal RNA (rRNA) -derived sRNA, SSU=small subunit, LSU=large subunit. Statistically significant findings are marked in bold.

**Table S10:** Twenty most significantly changed rDR species in HT users carried via HDL.

| HDL | Number | rDR | Annotation | log2 FoldChange | SE (log FoldChange) | Adjusted p-value |
|---|---|---|---|---|---|---|
| **HT user** | 1 | URS0000005270_9606 | Homo sapiens RNA, 5.8S ribosomal N3 | -0.083 | 0.529 | 0.997 |
| **POST vs. PRE** | 2 | URS000002B0D5_9606 | Homo sapiens 5S rRNA | 1.250 | 0.651 | 0.997 |
| | 3 | URS000003B35A_9606 | Homo sapiens 28S rRNA V8 region | 0.239 | 0.584 | 0.997 |
| | 4 | URS000005D9C4_9606 | Homo sapiens 28S rRNA V8 region | -0.154 | 0.464 | 0.997 |
| | 5 | URS0000086682_9606 | Homo sapiens 28S rRNA V8 region | -0.274 | 0.553 | 0.997 |
| | 6 | URS00000A5ABE_9606 | Homo sapiens 28S rRNA V8 region | -0.879 | 0.668 | 0.997 |
| | 7 | URS00000C39CA_9606 | Homo sapiens partial 28S rRNA | 0.018 | 0.693 | 0.997 |
| | 8 | URS00000D0A5B_9606 | Homo sapiens partial rRNA | -0.054 | 0.430 | 0.997 |
| | 9 | URS00000F9D45_9606 | Homo sapiens RNA, 5S ribosomal 12 | 1.078 | 0.821 | 0.997 |
| | 10 | URS000010CC88_9606 | Homo sapiens partial 18S rRNA | -1.321 | 0.909 | 0.997 |
| | 11 | URS00001152CA_9606 | Homo sapiens partial rRNA | -0.377 | 0.383 | 0.997 |
| | 12 | URS000013AA81_9606 | Homo sapiens 28S rRNA V8 region | -0.200 | 0.433 | 0.997 |
| | 13 | URS0000162C88_9606 | Homo sapiens rRNA LSU-S 5.8S | 0.078 | 0.514 | 0.997 |
| | 14 | URS0000165891_9606 | Homo sapiens SSU rRNA (18S) | -0.707 | 0.365 | 0.997 |
| | 15 | URS000016B819_9606 | Homo sapiens 28S rRNA V8 region | 0.027 | 0.477 | 0.997 |
| | 16 | URS0000171707_9606 | Homo sapiens partial rRNA | -0.322 | 0.462 | 0.997 |
| | 17 | URS000018FE5D_9606 | Homo sapiens LSU rRNA (28S) | -0.625 | 0.476 | 0.997 |
| | 18 | URS000019B9A7_9606 | Homo sapiens 28S rRNA V8 region | 0.544 | 0.479 | 0.997 |
| | 19 | URS00001AFAED_9606 | Homo sapiens partial rRNA | -0.657 | 0.563 | 0.997 |
| | 20 | URS00001C0ED1_9606 | Homo sapiens rRNA SSU 18S | 0.313 | 0.351 | 0.997 |
| **HT user** | 1 | URS000216BCCC_9606 | Homo sapiens (mitochondrial) LSU rRNA | 2.477 | 0.699 | 0.362 |
| **1h POST vs. PRE** | 2 | URS000068483A_9606 | Homo sapiens 5S rRNA | 2.755 | 0.760 | 0.362 |
| | 3 | URS0001A9A740_9606 | Homo sapiens SSU rRNA | 1.284 | 0.393 | 0.514 |
| | 4 | URS000216CA3F_9606 | Homo sapiens (mitochondrial) LSU rRNA | 2.146 | 0.658 | 0.514 |
| | 5 | URS0000684921_9606 | Homo sapiens 5S rRNA | 1.921 | 0.682 | 0.618 |
| | 6 | URS00006ACD21_9606 | Homo sapiens 5S rRNA (5S_rRNA) | 2.088 | 0.734 | 0.618 |
| | 7 | URS0000701637_9606 | Homo sapiens 5S rRNA | 1.791 | 0.689 | 0.618 |
| | 8 | URS00007125F9_9606 | Homo sapiens 5S rRNA | 1.661 | 0.706 | 0.618 |
| | 9 | URS0001A2E94F_9606 | Homo sapiens (mitochondrial) SSU rRNA | 1.625 | 0.630 | 0.618 |
| | 10 | URS0001A38BCF_9606 | Homo sapiens SSU rRNA | -1.021 | 0.426 | 0.618 |
| | 11 | URS0001A3E299_9606 | Homo sapiens (mitochondrial) SSU rRNA | 1.593 | 0.610 | 0.618 |
| | 12 | URS0001A48A19_9606 | Homo sapiens (mitochondrial) SSU rRNA | 1.650 | 0.605 | 0.618 |
| | 13 | URS0001A5DF53_9606 | Homo sapiens (mitochondrial) SSU rRNA | 1.727 | 0.733 | 0.618 |
| | 14 | URS0001A7336C_9606 | Homo sapiens (mitochondrial) SSU rRNA | 1.854 | 0.662 | 0.618 |

| | | | | | | |
|---|---|---|---|---|---|---|
| | 15 | URS0001A7C0B1_9606 | Homo sapiens (mitochondrial) SSU rRNA | 1.666 | 0.698 | 0.618 |
| | 16 | URS0001A7F20F_9606 | Homo sapiens SSU rRNA | 1.233 | 0.428 | 0.618 |
| | 17 | URS0001A83F53_9606 | Homo sapiens (mitochondrial) SSU rRNA | 2.025 | 0.689 | 0.618 |
| | 18 | URS0001A99078_9606 | Homo sapiens (mitochondrial) SSU rRNA | 1.696 | 0.703 | 0.618 |
| | 19 | URS0001AB3F32_9606 | Homo sapiens (mitochondrial) SSU rRNA | 1.607 | 0.642 | 0.618 |
| | 20 | URS0001ABB9D1_9606 | Homo sapiens (mitochondrial) SSU rRNA | 1.811 | 0.707 | 0.618 |
| **HT user** | 1 | URS0001A9A740_9606 | Homo sapiens SSU rRNA | 1.343 | 0.370 | 0.531 |
| **1h POST vs. POST** | 2 | URS0000005270_9606 | Homo sapiens RNA, 5.8S ribosomal N3 | 0.623 | 0.511 | 0.998 |
| | 3 | URS000002B0D5_9606 | Homo sapiens 5S rRNA | 0.136 | 0.587 | 0.998 |
| | 4 | URS000003B35A_9606 | Homo sapiens 28S rRNA V8 region | -0.243 | 0.575 | 0.998 |
| | 5 | URS000005D9C4_9606 | Homo sapiens 28S rRNA V8 region | 0.259 | 0.454 | 0.998 |
| | 6 | URS0000086682_9606 | Homo sapiens 28S rRNA V8 region | 0.561 | 0.533 | 0.998 |
| | 7 | URS00000A5ABE_9606 | Homo sapiens 28S rRNA V8 region | 0.516 | 0.655 | 0.998 |
| | 8 | URS00000C39CA_9606 | Homo sapiens partial 28S rRNA | -0.034 | 0.684 | 0.998 |
| | 9 | URS00000D0A5B_9606 | Homo sapiens partial rRNA | 0.224 | 0.404 | 0.998 |
| | 10 | URS00000F9D45_9606 | Homo sapiens RNA, 5S ribosomal 12 | 0.040 | 0.772 | 0.998 |
| | 11 | URS000010CC88_9606 | Homo sapiens partial 18S rRNA | 0.845 | 0.916 | 0.998 |
| | 12 | URS00001152CA_9606 | Homo sapiens partial rRNA | 0.627 | 0.365 | 0.998 |
| | 13 | URS000013AA81_9606 | Homo sapiens 28S rRNA V8 region | 0.054 | 0.438 | 0.998 |
| | 14 | URS0000162C88_9606 | Homo sapiens rRNA LSU-S 5.8S | 0.540 | 0.494 | 0.998 |
| | 15 | URS0000165891_9606 | Homo sapiens SSU rRNA (18S) | 0.453 | 0.359 | 0.998 |
| | 16 | URS000016B819_9606 | Homo sapiens 28S rRNA V8 region | -0.161 | 0.476 | 0.998 |
| | 17 | URS0000171707_9606 | Homo sapiens partial rRNA | 0.512 | 0.437 | 0.998 |
| | 18 | URS000018FE5D_9606 | Homo sapiens LSU rRNA (28S) | 0.226 | 0.455 | 0.998 |
| | 19 | URS000019B9A7_9606 | Homo sapiens 28S rRNA V8 region | -0.372 | 0.462 | 0.998 |
| | 20 | URS00001AFAED_9606 | Homo sapiens partial rRNA | 0.748 | 0.532 | 0.998 |

rDR= ribosomal RNA (rRNA) -derived sRNA, SSU=small subunit, LSU=large subunit.

**Table S11:** Twenty most significantly changed rDR species in nonusers carried via HDL.

| HDL | Number | rDR | Annotation | SE (log FoldChange) | Adjusted p-value |
|---|---|---|---|---|---|
| **Nonuser** | 1 | **URS0001A52223_9606** | Homo sapiens (mitochondrial) SSU rRNA | **0.619** | **0.005** |
| **POST vs. PRE** | 2 | **URS0000162C88_9606** | Homo sapiens rRNA LSU-S 5.8S | **0.356** | **0.017** |
| | 3 | **URS000080DE76_9606** | Homo sapiens 5.8S rRNA | **0.390** | **0.017** |
| | 4 | **URS00008C7B2C_9606** | Homo sapiens SSU rRNA | **0.358** | **0.017** |
| | 5 | **URS0001ABB9D1_9606** | Homo sapiens (mitochondrial) SSU rRNA | **0.543** | **0.017** |
| | 6 | **URS0001B1E32A_9606** | Homo sapiens SSU rRNA | **0.365** | **0.017** |
| | 7 | **URS0001B5B44A_9606** | Homo sapiens SSU rRNA | **0.357** | **0.017** |
| | 8 | **URS0001B77102_9606** | Homo sapiens (mitochondrial) SSU rRNA | **0.575** | **0.017** |
| | 9 | **URS0001BA83D1_9606** | Homo sapiens (mitochondrial) SSU rRNA | **0.740** | **0.017** |
| | 10 | **URS0002128349_9606** | Homo sapiens LSU rRNA | **0.338** | **0.017** |
| | 11 | **URS0002140352_9606** | Homo sapiens LSU rRNA | **0.329** | **0.017** |
| | 12 | **URS000214F71C_9606** | Homo sapiens LSU rRNA | **0.323** | **0.017** |
| | 13 | **URS00021ED94B_9606** | Homo sapiens 5.8S rRNA | **0.434** | **0.017** |
| | 14 | **URS0001A6CCD1_9606** | Homo sapiens (mitochondrial) SSU rRNA | **0.574** | **0.017** |
| | 15 | **URS0001B3F905_9606** | Homo sapiens (mitochondrial) SSU rRNA | **0.685** | **0.024** |
| | 16 | **URS0001A2E94F_9606** | Homo sapiens (mitochondrial) SSU rRNA | **0.727** | **0.029** |
| | 17 | **URS0001B28F7F_9606** | Homo sapiens (mitochondrial) SSU rRNA | **0.505** | **0.029** |
| | 18 | **URS0001B9C0C0_9606** | Homo sapiens SSU rRNA | **0.375** | **0.029** |
| | 19 | **URS000216BCCC_9606** | Homo sapiens (mitochondrial) LSU rRNA | **0.688** | **0.029** |
| | 20 | **URS000216CA3F_9606** | Homo sapiens (mitochondrial) LSU rRNA | **0.638** | **0.029** |
| **Nonuser** | 1 | **URS0001ABB9D1_9606** | Homo sapiens (mitochondrial) SSU rRNA | **0.537** | **0.000** |
| **1h POST vs. PRE** | 2 | **URS0001B28F7F_9606** | Homo sapiens (mitochondrial) SSU rRNA | **0.498** | **0.000** |
| | 3 | **URS0001A415C3_9606** | Homo sapiens (mitochondrial) SSU rRNA | **0.691** | **0.000** |
| | 4 | **URS0001A52223_9606** | Homo sapiens (mitochondrial) SSU rRNA | **0.615** | **0.000** |
| | 5 | **URS0001A53ABA_9606** | Homo sapiens (mitochondrial) SSU rRNA | **0.843** | **0.000** |
| | 6 | **URS0001A6CCD1_9606** | Homo sapiens (mitochondrial) SSU rRNA | **0.568** | **0.000** |
| | 7 | **URS0001BA83D1_9606** | Homo sapiens (mitochondrial) SSU rRNA | **0.735** | **0.000** |
| | 8 | **URS0001AB99B9_9606** | Homo sapiens (mitochondrial) SSU rRNA | **0.569** | **0.000** |
| | 9 | **URS0001B38803_9606** | Homo sapiens (mitochondrial) SSU rRNA | **0.896** | **0.000** |
| | 10 | **URS0001BA11BB_9606** | Homo sapiens (mitochondrial) SSU rRNA | **0.792** | **0.000** |
| | 11 | **URS000216BCCC_9606** | Homo sapiens (mitochondrial) LSU rRNA | **0.686** | **0.000** |
| | 12 | **URS000216CA3F_9606** | Homo sapiens (mitochondrial) LSU rRNA | **0.636** | **0.000** |
| | 13 | **URS0001A5136D_9606** | Homo sapiens (mitochondrial) SSU rRNA | **0.579** | **0.000** |
| | 14 | **URS0001B959E6_9606** | Homo sapiens (mitochondrial) SSU rRNA | **0.820** | **0.000** |
| | 15 | **URS0001BACCE4_9606** | Homo sapiens (mitochondrial) SSU rRNA | **0.550** | **0.000** |

| | | | | | |
|---|---|---|---|---|---|
| | 16 | **URS0001A94456_9606** | Homo sapiens (mitochondrial) SSU rRNA | **0.839** | **0.000** |
| | 17 | **URS0001ADA731_9606** | Homo sapiens (mitochondrial) SSU rRNA | **0.506** | **0.000** |
| | 18 | **URS0001B9B7CD_9606** | Homo sapiens (mitochondrial) SSU rRNA | **0.623** | **0.000** |
| | 19 | **URS0001AFD693_9606** | Homo sapiens (mitochondrial) SSU rRNA | **0.639** | **0.000** |
| | 20 | **URS0001A2E94F_9606** | Homo sapiens (mitochondrial) SSU rRNA | **0.720** | **0.000** |
| **Nonuser** | 1 | URS0000005270_9606 | Homo sapiens RNA, 5.8S ribosomal N3 | 0.439 | 0.997 |
| **1h POST vs. POST** | 2 | URS000002B0D5_9606 | Homo sapiens 5S rRNA | 0.595 | 0.997 |
| | 3 | URS000003B35A_9606 | Homo sapiens 28S rRNA V8 region | 0.517 | 0.997 |
| | 4 | URS000005D9C4_9606 | Homo sapiens 28S rRNA V8 region | 0.436 | 0.997 |
| | 5 | URS0000086682_9606 | Homo sapiens 28S rRNA V8 region | 0.545 | 0.997 |
| | 6 | URS00000A5ABE_9606 | Homo sapiens 28S rRNA V8 region | 0.457 | 0.997 |
| | 7 | URS00000C39CA_9606 | Homo sapiens partial 28S rRNA | 0.739 | 0.997 |
| | 8 | URS00000D0A5B_9606 | Homo sapiens partial rRNA | 0.351 | 0.997 |
| | 9 | URS00000F9D45_9606 | Homo sapiens RNA, 5S ribosomal 12 | 0.600 | 0.997 |
| | 10 | URS000010CC88_9606 | Homo sapiens partial 18S rRNA | 1.170 | 0.997 |
| | 11 | URS00001152CA_9606 | Homo sapiens partial rRNA | 0.364 | 0.997 |
| | 12 | URS000013AA81_9606 | Homo sapiens 28S rRNA V8 region | 0.482 | 0.997 |
| | 13 | URS0000162C88_9606 | Homo sapiens rRNA LSU-S 5.8S | 0.331 | 0.997 |
| | 14 | URS0000165891_9606 | Homo sapiens SSU rRNA (18S) | 0.373 | 0.997 |
| | 15 | URS000016B819_9606 | Homo sapiens 28S rRNA V8 region | 0.501 | 0.997 |
| | 16 | URS0000171707_9606 | Homo sapiens partial rRNA | 0.416 | 0.997 |
| | 17 | URS000018FE5D_9606 | Homo sapiens LSU rRNA (28S) | 0.336 | 0.997 |
| | 18 | URS000019B9A7_9606 | Homo sapiens 28S rRNA V8 region | 0.447 | 0.997 |
| | 19 | URS00001AFAED_9606 | Homo sapiens partial rRNA | 0.307 | 0.997 |
| | 20 | URS00001C0ED1_9606 | Homo sapiens rRNA SSU 18S | 0.333 | 0.997 |

rDR= ribosomal RNA (rRNA) -derived sRNA, SSU=small subunit, LSU=large subunit. Statistically significant findings are marked in bold.

**Table S12:** Twenty most significantly changed tDR species in HT users carried via EVs.

| EV | Number | tDR | tRNA | Anticodon | log2 FoldChange | SE (log FoldChange) | Adjusted p-value |
|---|---|---|---|---|---|---|---|
| **HT user** | 1 | URS000009AC8B_9606 | tRNA-Gly | (GCC) | 0.061 | 0.194 | 0.989 |
| **POST vs. PRE** | 2 | URS000009DDCA_9606 | tRNA-Val | (CAC) | -0.268 | 0.483 | 0.989 |
| | 3 | URS0000120E41_9606 | tRNA-Leu | (AAG) | -0.361 | 0.923 | 0.989 |
| | 4 | URS000013899F_9606 | tRNA-Asp | (GTC) | -0.404 | 0.460 | 0.989 |
| | 5 | URS000013B42D_9606 | tRNA-Gly | (GCC) | 0.026 | 0.155 | 0.989 |
| | 6 | URS00001AD596_9606 | tRNA-Lys | (CTT) | -0.383 | 0.288 | 0.989 |
| | 7 | URS00001B506A_9606 | tRNA-Leu | (TAG) | 0.276 | 0.844 | 0.989 |
| | 8 | URS00002064F6_9606 | tRNA-Val | (CAC) | -0.069 | 0.184 | 0.989 |
| | 9 | URS000022DD4A_9606 | tRNA-Lys | (TTT) | 0.308 | 0.422 | 0.989 |
| | 10 | URS0000287398_9606 | tRNA-Glu | (TTC) | 0.078 | 0.538 | 0.989 |
| | 11 | URS00002B4CE5_9606 | tRNA-Val | (CAC) | 0.255 | 0.256 | 0.989 |
| | 12 | URS000038803E_9606 | tRNA-Val | (TAC) | 0.697 | 1.051 | 0.989 |
| | 13 | URS00003C9A26_9606 | tRNA-Glu | (TTC) | 0.289 | 0.281 | 0.989 |
| | 14 | URS00004310BE_9606 | tRNA-Val | (AAC) | 0.144 | 0.160 | 0.989 |
| | 15 | URS000044BAE3_9606 | tRNA-His | (GTG) | 0.325 | 0.572 | 0.989 |
| | 16 | URS00004BF687_9606 | tRNA-Gly | (CCC) | -0.142 | 0.195 | 0.989 |
| | 17 | URS00004D9E92_9606 | tRNA-Lys | (CTT) | -0.393 | 0.259 | 0.989 |
| | 18 | URS00004F0321_9606 | tRNA-Glu | (CTC) | 0.027 | 0.200 | 0.989 |
| | 19 | URS0000502C74_9606 | tRNA-Gly | (CCC) | 0.245 | 0.231 | 0.989 |
| | 20 | URS00006174C2_9606 | tRNA-Asp | (GTC) | -0.223 | 0.268 | 0.989 |
| **HT user** | 1 | URS000064506B_9606 | tRNA-Leu | (AAG) | -3.337 | 1.114 | 0.165 |
| **1h POST vs. PRE** | 2 | URS000070F1D2_9606 | tRNA-Gln | (TTG) | 3.426 | 1.103 | 0.165 |
| | 3 | URS000047CD44_9606 | tRNA-Arg | (TCT) | -1.769 | 0.606 | 0.165 |
| | 4 | URS00000AED6F_9606 | tRNA-Gly | (TCC) | -2.748 | 1.001 | 0.213 |
| | 5 | URS0000120E41_9606 | tRNA-Leu | (AAG) | -1.636 | 0.962 | 0.677 |
| | 6 | URS000038803E_9606 | tRNA-Val | (TAC) | 1.914 | 1.020 | 0.677 |
| | 7 | URS0000679FAF_9606 | tRNA-Lys | (CTT) | 1.016 | 0.535 | 0.677 |
| | 8 | URS000071ED2F_9606 | tRNA-Val | (AAC) | 1.443 | 0.766 | 0.677 |
| | 9 | URS000072BBEB_9606 | tRNA-Glu | (TCC) | 1.076 | 0.593 | 0.677 |
| | 10 | URS00000C18F2_9606 | tRNA-Pro | (CGG) | -1.197 | 0.566 | 0.677 |
| | 11 | URS0000493225_9606 | tRNA-Pro | (TGG) | 1.682 | 0.854 | 0.677 |
| | 12 | URS0000572B72_9606 | tRNA-Leu | (CAA) | -1.241 | 0.700 | 0.677 |
| | 13 | URS000059900F_9606 | tRNA-Arg | (CCG) | 2.745 | 1.471 | 0.677 |
| | 14 | URS0000677B31_9606 | tRNA-Ala | (AGC) | 1.538 | 0.911 | 0.677 |

| | | | | | | | |
|---|---|---|---|---|---|---|---|
| | 15 | URS0000758799_9606 | tRNA-Tyr | (GTA) | 2.029 | 1.181 | 0.677 |
| | 16 | URS00006C8EDF_9606 | tRNA-Ala | (TGC) | -1.371 | 0.784 | 0.677 |
| | 17 | URS0000092302_9606 | tRNA-Ser | (TGA) | -2.118 | 0.993 | 0.677 |
| | 18 | URS000064E10F_9606 | tRNA-Leu | (TAA) | -2.023 | 1.091 | 0.677 |
| | 19 | URS0000716B70_9606 | tRNA-Gly | (TCC) | -1.566 | 0.885 | 0.677 |
| | 20 | URS00006BE24E_9606 | tRNA-Asn | (GTT) | -1.829 | 1.122 | 0.727 |
| **HT user** | 1 | **URS000070F1D2_9606** | tRNA-Gln | (TTG) | 4.404 | 1.111 | **0.010** |
| **1h POST vs. POST** | 2 | URS0000679FAF_9606 | tRNA-Lys | (CTT) | 1.570 | 0.541 | 0.151 |
| | 3 | URS000059900F_9606 | tRNA-Arg | (CCG) | 4.269 | 1.494 | 0.151 |
| | 4 | URS00006D74B2_9606 | tRNA-Asp | (GTC) | 1.048 | 0.346 | 0.151 |
| | 5 | URS000072A930_9606 | tRNA-Lys | (TTT) | 2.254 | 0.818 | 0.165 |
| | 6 | URS00003C9A26_9606 | tRNA-Glu | (TTC) | -0.720 | 0.283 | 0.196 |
| | 7 | URS000064506B_9606 | tRNA-Leu | (AAG) | -2.801 | 1.122 | 0.196 |
| | 8 | URS00006C8EDF_9606 | tRNA-Ala | (TGC) | -2.013 | 0.772 | 0.196 |
| | 9 | URS0000092302_9606 | tRNA-Ser | (TGA) | -2.491 | 0.985 | 0.196 |
| | 10 | URS000064E10F_9606 | tRNA-Leu | (TAA) | -2.392 | 1.084 | 0.385 |
| | 11 | URS000047CD44_9606 | tRNA-Arg | (TCT) | -1.316 | 0.612 | 0.404 |
| | 12 | URS00002D40C8_9606 | tRNA-Pro | (TGG) | 1.317 | 0.650 | 0.463 |
| | 13 | URS0000493225_9606 | tRNA-Pro | (TGG) | 1.741 | 0.858 | 0.463 |
| | 14 | URS000037D0FB_9606 | tRNA-Ser | (TGA) | -1.189 | 0.628 | 0.587 |
| | 15 | URS0000209048_9606 | tRNA-Ala | (AGC) | -1.422 | 0.771 | 0.611 |
| | 16 | URS00006174C2_9606 | tRNA-Asp | (GTC) | 0.466 | 0.267 | 0.668 |
| | 17 | URS00000AED6F_9606 | tRNA-Gly | (TCC) | -1.796 | 1.012 | 0.668 |
| | 18 | URS0000572B72_9606 | tRNA-Leu | (CAA) | -1.164 | 0.679 | 0.678 |
| | 19 | URS000072BBEB_9606 | tRNA-Glu | (TCC) | 0.943 | 0.584 | 0.784 |
| | 20 | URS000062C4DE_9606 | tRNA-Pro | (TGG) | -2.089 | 1.352 | 0.784 |

tDR= transfer RNA (tRNA) -derived sRNA. Statistically significant findings are marked in bold.

**Table S13:** Twenty most significantly changed tDR species in nonusers carried via EVs.

| EV | Number | tDR | tRNA | Anticodon | log2 FoldChange | SE (log FoldChange) | Adjusted p-value |
|---|---|---|---|---|---|---|---|
| **Nonuser POST vs. PRE** | 1 | URS0000738059_9606 | tRNA-Arg | (TCT) | 4.453 | 1.406 | 0.217 |
| | 2 | URS000009AC8B_9606 | tRNA-Gly | (GCC) | 0.085 | 0.210 | 0.994 |
| | 3 | URS000009DDCA_9606 | tRNA-Val | (CAC) | -0.853 | 0.897 | 0.994 |
| | 4 | URS0000120E41_9606 | tRNA-Leu | (AAG) | -0.115 | 0.762 | 0.994 |
| | 5 | URS000013899F_9606 | tRNA-Asp | (GTC) | -1.257 | 0.845 | 0.994 |
| | 6 | URS0000161979_9606 | tRNA-Val | (AAC) | -1.025 | 0.573 | 0.994 |
| | 7 | URS00001AD596_9606 | tRNA-Lys | (CTT) | -0.870 | 0.589 | 0.994 |
| | 8 | URS00001B506A_9606 | tRNA-Leu | (TAG) | 0.210 | 0.768 | 0.994 |
| | 9 | URS00002064F6_9606 | tRNA-Val | (CAC) | -0.501 | 0.561 | 0.994 |
| | 10 | URS0000287398_9606 | tRNA-Glu | (TTC) | 0.810 | 0.539 | 0.994 |
| | 11 | URS00002B4CE5_9606 | tRNA-Val | (CAC) | 0.300 | 0.711 | 0.994 |
| | 12 | URS00003C9A26_9606 | tRNA-Glu | (TTC) | -0.773 | 0.626 | 0.994 |
| | 13 | URS00004310BE_9606 | tRNA-Val | (AAC) | -0.371 | 0.454 | 0.994 |
| | 14 | URS00004BF687_9606 | tRNA-Gly | (CCC) | -0.109 | 0.268 | 0.994 |
| | 15 | URS00004D9E92_9606 | tRNA-Lys | (CTT) | -0.866 | 0.596 | 0.994 |
| | 16 | URS00004F0321_9606 | tRNA-Glu | (CTC) | 0.041 | 0.320 | 0.994 |
| | 17 | URS0000502C74_9606 | tRNA-Gly | (CCC) | -0.783 | 0.406 | 0.994 |
| | 18 | URS00006174C2_9606 | tRNA-Asp | (GTC) | -0.335 | 0.347 | 0.994 |
| | 19 | URS000061D582_9606 | tRNA-Val | (AAC) | -0.477 | 0.542 | 0.994 |
| | 20 | URS0000630B8A_9606 | tRNA-Leu | (AAG) | 0.390 | 0.762 | 0.994 |
| **Nonuser 1h POST vs. PRE** | 1 | URS000064506B_9606 | tRNA-Leu | (AAG) | -3.113 | 1.048 | 0.290 |
| | 2 | URS000072A930_9606 | tRNA-Lys | (TTT) | -3.161 | 1.102 | 0.290 |
| | 3 | URS000009AC8B_9606 | tRNA-Gly | (GCC) | -0.044 | 0.213 | 0.989 |
| | 4 | URS000009DDCA_9606 | tRNA-Val | (CAC) | -0.049 | 0.897 | 0.989 |
| | 5 | URS0000120E41_9606 | tRNA-Leu | (AAG) | -0.812 | 0.811 | 0.989 |
| | 6 | URS000013899F_9606 | tRNA-Asp | (GTC) | -0.566 | 0.846 | 0.989 |
| | 7 | URS000013B42D_9606 | tRNA-Gly | (GCC) | -0.196 | 0.236 | 0.989 |
| | 8 | URS0000161979_9606 | tRNA-Val | (AAC) | -0.486 | 0.574 | 0.989 |
| | 9 | URS00001AD596_9606 | tRNA-Lys | (CTT) | -0.442 | 0.590 | 0.989 |
| | 10 | URS00001B506A_9606 | tRNA-Leu | (TAG) | 0.286 | 0.781 | 0.989 |
| | 11 | URS00002064F6_9606 | tRNA-Val | (CAC) | 0.247 | 0.560 | 0.989 |
| | 12 | URS000022DD4A_9606 | tRNA-Lys | (TTT) | -0.724 | 0.496 | 0.989 |
| | 13 | URS0000287398_9606 | tRNA-Glu | (TTC) | 0.008 | 0.564 | 0.989 |
| | 14 | URS00002B4CE5_9606 | tRNA-Val | (CAC) | 1.018 | 0.709 | 0.989 |
| | 15 | URS000038803E_9606 | tRNA-Val | (TAC) | -1.909 | 0.994 | 0.989 |

| | | | | | | | |
|---|---|---|---|---|---|---|---|
| | 16 | URS00003C9A26_9606 | tRNA-Glu | (TTC) | -0.840 | 0.628 | 0.989 |
| | 17 | URS00004310BE_9606 | tRNA-Val | (AAC) | 0.040 | 0.454 | 0.989 |
| | 18 | URS000044BAE3_9606 | tRNA-His | (GTG) | 0.795 | 0.909 | 0.989 |
| | 19 | URS00004BF687_9606 | tRNA-Gly | (CCC) | -0.072 | 0.272 | 0.989 |
| | 20 | URS00004D9E92_9606 | tRNA-Lys | (CTT) | -0.541 | 0.597 | 0.989 |
| **Nonuser** | 1 | URS000072A930_9606 | tRNA-Lys | (TTT) | -3.842 | 1.085 | 0.056 |
| **1h POST vs. POST** | 2 | URS0000679FAF_9606 | tRNA-Lys | (CTT) | -2.421 | 0.804 | 0.122 |
| | 3 | URS000071ED2F_9606 | tRNA-Val | (AAC) | 2.567 | 0.828 | 0.122 |
| | 4 | URS000064506B_9606 | tRNA-Leu | (AAG) | -2.773 | 1.031 | 0.253 |
| | 5 | URS0000738059_9606 | tRNA-Arg | (TCT) | -3.552 | 1.379 | 0.283 |
| | 6 | URS000034AAC2_9606 | tRNA-Gly | (TCC) | 2.114 | 0.858 | 0.323 |
| | 7 | URS0000209048_9606 | tRNA-Ala | (AGC) | 1.664 | 0.693 | 0.328 |
| | 8 | URS00006C6D0A_9606 | tRNA-Ala | (TGC) | 1.906 | 0.893 | 0.579 |
| | 9 | URS000038803E_9606 | tRNA-Val | (TAC) | -1.904 | 0.975 | 0.660 |
| | 10 | URS000064074C_9606 | tRNA-Glu | (TTC) | 1.259 | 0.630 | 0.660 |
| | 11 | URS0000748C4B_9606 | tRNA-Val | (CAC) | 1.061 | 0.545 | 0.660 |
| | 12 | URS000030BAD5_9606 | tRNA-iMet | (CAT) | 1.873 | 1.000 | 0.718 |
| | 13 | URS000029CCC5_9606 | tRNA-Phe | (GAA) | -2.589 | 1.436 | 0.755 |
| | 14 | URS00007047C1_9606 | tRNA suppressor | (TTA) | 1.999 | 1.123 | 0.755 |
| | 15 | URS000009DDCA_9606 | tRNA-Val | (CAC) | 0.804 | 0.894 | 0.804 |
| | 16 | URS0000120E41_9606 | tRNA-Leu | (AAG) | -0.696 | 0.784 | 0.804 |
| | 17 | URS000013899F_9606 | tRNA-Asp | (GTC) | 0.691 | 0.840 | 0.804 |
| | 18 | URS000013B42D_9606 | tRNA-Gly | (GCC) | -0.188 | 0.228 | 0.804 |
| | 19 | URS0000161979_9606 | tRNA-Val | (AAC) | 0.539 | 0.572 | 0.804 |
| | 20 | URS00002064F6_9606 | tRNA-Val | (CAC) | 0.747 | 0.555 | 0.804 |

tDR= transfer RNA (tRNA) -derived sRNA.

**Table S14:** Twenty most significantly changed tDR species in HT users carried via HDL.

| HDL | Number | tDR | | | log2 FoldChange | SE (log FoldChange) | Adjusted p-value |
|---|---|---|---|---|---|---|---|
| **HT user POST vs. PRE** | 1 | URS000072BBEB_9606 | tRNA-Glu | (TCC) | 2.972 | 0.817 | 0.074 |
| | 2 | URS0000639DBE_9606 | tRNA-Glu | (TTC) | 1.200 | 0.394 | 0.310 |
| | 3 | URS00001B506A_9606 | tRNA-Leu | (TAG) | 4.380 | 1.544 | 0.408 |
| | 4 | URS00004F0321_9606 | tRNA-Glu | (CTC) | 0.607 | 0.255 | 0.660 |
| | 5 | URS00005A1E0D_9606 | tRNA-Asn | (GTT) | -2.863 | 1.166 | 0.660 |
| | 6 | URS0000635088_9606 | tRNA-Glu | (CTC) | 0.646 | 0.271 | 0.660 |
| | 7 | URS0000732902_9606 | tRNA-Tyr | (GTA) | -1.967 | 0.779 | 0.660 |
| | 8 | URS0000755767_9606 | tRNA-Tyr | (GTA) | -2.933 | 1.303 | 0.765 |
| | 9 | URS000064BBA5_9606 | tRNA-Phe | (GAA) | -2.646 | 1.185 | 0.765 |
| | 10 | URS000005AEAB_9606 | tRNA-Leu | (TAG) | 0.328 | 1.816 | 1.000 |
| | 11 | URS000009AC8B_9606 | tRNA-Gly | (GCC) | 0.057 | 0.182 | 1.000 |
| | 12 | URS000009DDCA_9606 | tRNA-Val | (CAC) | 0.335 | 0.653 | 1.000 |
| | 13 | URS00000AED6F_9606 | tRNA-Gly | (TCC) | -0.160 | 1.821 | 1.000 |
| | 14 | URS00000C18F2_9606 | tRNA-Pro | (CGG) | -0.957 | 0.832 | 1.000 |
| | 15 | URS00000F30A4_9606 | tRNA-Thr | (TGT) | 0.165 | 3.687 | 1.000 |
| | 16 | URS00000FB60D_9606 | tRNA-Leu | (CAG) | 2.439 | 2.107 | 1.000 |
| | 17 | URS0000120E41_9606 | tRNA-Leu | (AAG) | 2.389 | 1.995 | 1.000 |
| | 18 | URS0000121433_9606 | tRNA-iMet | (CAT) | 0.684 | 1.443 | 1.000 |
| | 19 | URS000013899F_9606 | tRNA-Asp | (GTC) | 0.920 | 0.862 | 1.000 |
| | 20 | URS000013B42D_9606 | tRNA-Gly | (GCC) | 0.026 | 0.153 | 1.000 |
| **HT user 1h POST vs. PRE** | 1 | **URS0000639DBE_9606** | tRNA-Glu | (TTC) | 1.448 | 0.394 | **0.050** |
| | 2 | **URS000072BBEB_9606** | tRNA-Glu | (TTC) | 2.901 | 0.815 | **0.050** |
| | 3 | URS000005AEAB_9606 | tRNA-Leu | (TAG) | 0.916 | 1.808 | 1.000 |
| | 4 | URS000009AC8B_9606 | tRNA-Gly | (GCC) | 0.029 | 0.181 | 1.000 |
| | 5 | URS000009DDCA_9606 | tRNA-Val | (CAC) | 0.191 | 0.651 | 1.000 |
| | 6 | URS00000AED6F_9606 | tRNA-Gly | (TCC) | -1.469 | 1.871 | 1.000 |
| | 7 | URS00000C18F2_9606 | tRNA-Pro | (CGG) | -2.048 | 0.847 | 1.000 |
| | 8 | URS00000F30A4_9606 | tRNA-Thr | (TGT) | 0.000 | 3.698 | 1.000 |
| | 9 | URS00000FB60D_9606 | tRNA-Leu | (CAG) | 2.151 | 2.112 | 1.000 |
| | 10 | URS0000120E41_9606 | tRNA-Leu | (AAG) | 2.010 | 1.997 | 1.000 |
| | 11 | URS0000121433_9606 | tRNA-iMet | (CAT) | 1.296 | 1.425 | 1.000 |
| | 12 | URS000013899F_9606 | tRNA-Asp | (GTC) | 0.586 | 0.860 | 1.000 |
| | 13 | URS000013B42D_9606 | tRNA-Gly | (GCC) | -0.099 | 0.154 | 1.000 |
| | 14 | URS0000145C5E_9606 | tRNA-Met | (CAT) | 0.483 | 3.652 | 1.000 |

| | | | | | | | |
|---|---|---|---|---|---|---|---|
| | 15 | URS00001618FC_9606 | tRNA-Trp | (CCA) | 0.016 | 3.691 | 1.000 |
| | 16 | URS0000161979_9606 | tRNA-Val | (AAC) | 0.243 | 0.479 | 1.000 |
| | 17 | URS00001A86BB_9606 | tRNA-Val | (CAC) | 0.026 | 0.848 | 1.000 |
| | 18 | URS00001AD596_9606 | tRNA-Lys | (CTT) | 0.261 | 0.622 | 1.000 |
| | 19 | URS00001B506A_9606 | tRNA-Leu | (TAG) | 3.587 | 1.562 | 1.000 |
| | 20 | URS00001D909A_9606 | tRNA-Arg | (CCG) | 0.250 | 3.661 | 1.000 |
| **HT user** | 1 | URS0000732902_9606 | tRNA-Tyr | (GTA) | 2.305 | 0.757 | 0.368 |
| **1h POST vs. POST** | 2 | URS0000209048_9606 | tRNA-Ala | (AGC) | -2.571 | 0.858 | 0.368 |
| | 3 | URS000005AEAB_9606 | tRNA-Leu | (TAG) | 0.589 | 1.794 | 1.000 |
| | 4 | URS000009AC8B_9606 | tRNA-Gly | (GCC) | -0.028 | 0.178 | 1.000 |
| | 5 | URS000009DDCA_9606 | tRNA-Val | (CAC) | -0.143 | 0.645 | 1.000 |
| | 6 | URS00000AED6F_9606 | tRNA-Gly | (TCC) | -1.309 | 1.858 | 1.000 |
| | 7 | URS00000C18F2_9606 | tRNA-Pro | (CGG) | -1.091 | 0.855 | 1.000 |
| | 8 | URS00000F30A4_9606 | tRNA-Thr | (TGT) | -0.707 | 3.687 | 1.000 |
| | 9 | URS00000FB60D_9606 | tRNA-Leu | (CAG) | -0.287 | 2.023 | 1.000 |
| | 10 | URS0000120E41_9606 | tRNA-Leu | (AAG) | -0.379 | 1.915 | 1.000 |
| | 11 | URS0000121433_9606 | tRNA-iMet | (CAT) | 0.612 | 1.388 | 1.000 |
| | 12 | URS000013899F_9606 | tRNA-Asp | (GTC) | -0.334 | 0.851 | 1.000 |
| | 13 | URS000013B42D_9606 | tRNA-Gly | (GCC) | -0.125 | 0.151 | 1.000 |
| | 14 | URS0000145C5E_9606 | tRNA-Met | (CAT) | 1.152 | 3.649 | 1.000 |
| | 15 | URS00001618FC_9606 | tRNA-Trp | (CCA) | -0.370 | 3.673 | 1.000 |
| | 16 | URS0000161979_9606 | tRNA-Val | (AAC) | -0.239 | 0.474 | 1.000 |
| | 17 | URS00001A86BB_9606 | tRNA-Val | (CAC) | 0.348 | 0.837 | 1.000 |
| | 18 | URS00001AD596_9606 | tRNA-Lys | (CTT) | 0.623 | 0.622 | 1.000 |
| | 19 | URS00001B506A_9606 | tRNA-Leu | (TAG) | -0.793 | 1.403 | 1.000 |
| | 20 | URS00001D909A_9606 | tRNA-Arg | (CCG) | -0.427 | 3.643 | 1.000 |

tDR= transfer RNA (tRNA) -derived sRNA. Statistically significant findings are marked in bold.

**Table S15:** Twenty most significantly changed tDR species in nonusers carried via HDL.

| HDL | Number | tDR | | | log2 FoldChange | SE (log FoldChange) | Adjusted p-value |
|---|---|---|---|---|---|---|---|
| **Nonuser** | 1 | **URS00006B479B_9606** | tRNA-Gly | (CCC) | -5.086 | 1.199 | **0.003** |
| **POST vs. PRE** | 2 | **URS00007131F2_9606** | tRNA-Gly | (CCC) | -5.041 | 1.328 | **0.010** |
| | 3 | **URS000013899F_9606** | tRNA-Asp | (GTC) | -1.570 | 0.482 | **0.037** |
| | 4 | **URS0000493225_9606** | tRNA-Pro | (TGG) | -3.122 | 0.957 | **0.037** |
| | 5 | **URS00003C9A26_9606** | tRNA-Glu | (TTC) | 1.382 | 0.442 | **0.045** |
| | 6 | **URS00006D74B2_9606** | tRNA-Asp | (GTC) | -2.251 | 0.730 | **0.045** |
| | 7 | **URS00001AD596_9606** | tRNA-Lys | (CTT) | -1.639 | 0.546 | **0.048** |
| | 8 | **URS000069E2A5_9606** | tRNA-Ala | (TGC) | -3.642 | 1.253 | **0.048** |
| | 9 | **URS00002034DC_9606** | tRNA-Ser | (GCT) | -4.832 | 1.655 | **0.048** |
| | 10 | **URS0000664EAD_9606** | tRNA-Ser | (GCT) | -4.839 | 1.665 | **0.048** |
| | 11 | **URS00004CF258_9606** | tRNA-Ser | (GCT) | -4.726 | 1.644 | **0.048** |
| | 12 | URS000071ED2F_9606 | tRNA-Val | (AAC) | -2.343 | 0.871 | 0.072 |
| | 13 | URS000067AC87_9606 | tRNA-Ser | (GCT) | -4.634 | 1.722 | 0.072 |
| | 14 | URS00000C18F2_9606 | tRNA-Pro | (CGG) | -2.524 | 0.998 | 0.097 |
| | 15 | URS00002D40C8_9606 | tRNA-Pro | (TGG) | -2.296 | 0.912 | 0.097 |
| | 16 | URS00001A72CE_9606 | tRNA-Trp | (CCA) | -4.548 | 1.779 | 0.097 |
| | 17 | URS0000659544_9606 | tRNA-Lys | (CTT) | -1.301 | 0.526 | 0.103 |
| | 18 | URS00005B30A9_9606 | tRNA-Ala | (CGC) | -2.371 | 0.983 | 0.115 |
| | 19 | URS000047EBB5_9606 | tRNA-Ser | (GCT) | -4.231 | 1.795 | 0.127 |
| | 20 | URS000064A807_9606 | tRNA-Asp | (GTC) | -3.720 | 1.601 | 0.132 |
| **Nonuser** | 1 | **URS00003C9A26_9606** | tRNA-Glu | (TTC) | 1.726 | 0.442 | **0.025** |
| **1h POST vs. PRE** | 2 | URS0000639DBE_9606 | tRNA-Glu | (TTC) | 1.449 | 0.508 | 0.574 |
| | 3 | URS00002B4CE5_9606 | tRNA-Val | (CAC) | -1.175 | 0.451 | 0.809 |
| | 4 | URS0000493225_9606 | tRNA-Pro | (TGG) | -2.320 | 0.937 | 0.877 |
| | 5 | URS00000C18F2_9606 | tRNA-Pro | (CGG) | -2.393 | 1.006 | 0.917 |
| | 6 | URS000052A1C9_9606 | tRNA-Ala | (AGC) | -5.251 | 2.288 | 0.956 |
| | 7 | URS000005AEAB_9606 | tRNA-Leu | (TAG) | 0.937 | 1.446 | 1.000 |
| | 8 | URS000009AC8B_9606 | tRNA-Gly | (GCC) | -0.013 | 0.230 | 1.000 |
| | 9 | URS000009DDCA_9606 | tRNA-Val | (CAC) | -0.428 | 0.622 | 1.000 |
| | 10 | URS00000AED6F_9606 | tRNA-Gly | (TCC) | 0.086 | 1.799 | 1.000 |
| | 11 | URS00000F30A4_9606 | tRNA-Thr | (TGT) | 0.236 | 3.661 | 1.000 |
| | 12 | URS00000FB60D_9606 | tRNA-Leu | (CAG) | 1.011 | 1.422 | 1.000 |
| | 13 | URS0000120E41_9606 | tRNA-Leu | (AAG) | 1.532 | 1.648 | 1.000 |
| | 14 | URS0000121433_9606 | tRNA-iMet | (CAT) | -1.284 | 1.279 | 1.000 |
| | 15 | URS000013899F_9606 | tRNA-Asp | (GTC) | -0.544 | 0.473 | 1.000 |

| | | | | | | | |
|---|---|---|---|---|---|---|---|
| | 16 | URS000013B42D_9606 | tRNA-Gly | (GCC) | 0.139 | 0.206 | 1.000 |
| | 17 | URS0000145C5E_9606 | tRNA-Met | (CAT) | -0.915 | 1.555 | 1.000 |
| | 18 | URS00001618FC_9606 | tRNA-Trp | (CCA) | -3.232 | 3.631 | 1.000 |
| | 19 | URS0000161979_9606 | tRNA-Val | (AAC) | 0.806 | 0.534 | 1.000 |
| | 20 | URS00001A86BB_9606 | tRNA-Val | (CAC) | 0.247 | 1.323 | 1.000 |
| **Nonuser** | 1 | URS000071ED2F_9606 | tRNA-Val | (AAC) | 2.620 | 0.825 | 0.395 |
| **1h POST vs. POST** | 2 | URS00001AD596_9606 | tRNA-Lys | (CTT) | 1.541 | 0.543 | 0.404 |
| | 3 | URS00007131F2_9606 | tRNA-Gly | (CCC) | 3.728 | 1.314 | 0.404 |
| | 4 | URS00002B4CE5_9606 | tRNA-Val | (CAC) | -1.143 | 0.439 | 0.483 |
| | 5 | URS00006B479B_9606 | tRNA-Gly | (CCC) | 3.195 | 1.194 | 0.483 |
| | 6 | URS00005B30A9_9606 | tRNA-Ala | (CGC) | 2.316 | 0.970 | 0.746 |
| | 7 | URS0000702883_9606 | tRNA-Lys | (TTT) | 2.079 | 0.900 | 0.789 |
| | 8 | URS000013899F_9606 | tRNA-Asp | (GTC) | 1.026 | 0.475 | 0.814 |
| | 9 | URS0000333F2E_9606 | tRNA-Tyr | (GTA) | 3.411 | 1.580 | 0.814 |
| | 10 | URS00004D9E92_9606 | tRNA-Lys | (CTT) | 1.231 | 0.548 | 0.814 |
| | 11 | URS000005AEAB_9606 | tRNA-Leu | (TAG) | 1.982 | 1.402 | 1.000 |
| | 12 | URS000009AC8B_9606 | tRNA-Gly | (GCC) | -0.051 | 0.226 | 1.000 |
| | 13 | URS000009DDCA_9606 | tRNA-Val | (CAC) | -0.573 | 0.612 | 1.000 |
| | 14 | URS00000AED6F_9606 | tRNA-Gly | (TCC) | 1.910 | 1.789 | 1.000 |
| | 15 | URS00000C18F2_9606 | tRNA-Pro | (CGG) | 0.131 | 1.022 | 1.000 |
| | 16 | URS00000F30A4_9606 | tRNA-Thr | (TGT) | -0.616 | 3.623 | 1.000 |
| | 17 | URS00000FB60D_9606 | tRNA-Leu | (CAG) | 1.740 | 1.373 | 1.000 |
| | 18 | URS0000120E41_9606 | tRNA-Leu | (AAG) | 1.427 | 1.600 | 1.000 |
| | 19 | URS0000121433_9606 | tRNA-iMet | (CAT) | 0.168 | 1.263 | 1.000 |
| | 20 | URS000013B42D_9606 | tRNA-Gly | (GCC) | 0.046 | 0.201 | 1.000 |

tDR= transfer RNA (tRNA) -derived sRNA. Statistically significant findings are marked in bold.

**Table S16**. KEGG pathways of differentially expressed miRs in EV and HDL particles.

| | KEGG pathway | Number of regulating miRs | Number of regulated genes | p-value |
|---|---|---|---|---|
| **EV miRs** | TGF-beta signaling pathway | 5 | 28 | 0.000 |
| | Signaling pathways regulating pluripotency of stem cells | 5 | 46 | 0.000 |
| | Hippo signaling pathway | 5 | 35 | 0.000 |
| | Proteoglycans in cancer | 4 | 49 | 0.001 |
| | Phosphatidylinositol signaling system | 5 | 27 | 0.001 |
| | Melanoma | 4 | 23 | 0.003 |
| | Glioma | 4 | 19 | 0.005 |
| | mTOR signaling pathway | 4 | 22 | 0.008 |
| | Regulation of actin cytoskeleton | 4 | 55 | 0.008 |
| | Transcriptional misregulation in cancer | 5 | 43 | 0.008 |
| | Thyroid hormone signaling pathway | 5 | 28 | 0.008 |
| | Thyroid hormone synthesis | 4 | 16 | 0.009 |
| | Dorso-ventral axis formation | 4 | 12 | 0.012 |
| | Pancreatic cancer | 4 | 21 | 0.012 |
| | Adherens junction | 5 | 23 | 0.013 |
| | T cell receptor signaling pathway | 4 | 30 | 0.015 |
| | FoxO signaling pathway | 4 | 34 | 0.015 |
| | Small cell lung cancer | 4 | 25 | 0.020 |
| | Non-small cell lung cancer | 4 | 16 | 0.021 |
| | Protein processing in endoplasmic reticulum | 3 | 42 | 0.029 |
| | Circadian rhythm | 2 | 12 | 0.031 |
| | Inositol phosphate metabolism | 5 | 17 | 0.031 |
| | Neurotrophin signaling pathway | 5 | 30 | 0.031 |
| | Focal adhesion | 4 | 50 | 0.033 |
| | PI3K-Akt signaling pathway | 4 | 76 | 0.034 |
| | Ras signaling pathway | 4 | 52 | 0.035 |
| | TNF signaling pathway | 4 | 29 | 0.041 |
| | Pathways in cancer | 4 | 87 | 0.048 |
| **HDL miRs** | Glycosphingolipid biosynthesis - lacto and neolacto series | 3 | 5 | 0.000 |
| | ECM-receptor interaction | 3 | 16 | 0.000 |
| | Proteoglycans in cancer | 3 | 34 | 0.005 |

| | | | | |
|---|---|---|---|---|
| | Dorso-ventral axis formation | 3 | 10 | 0.025 |
| | Transcriptional misregulation in cancer | 3 | 36 | 0.025 |
| | Amoebiasis | 3 | 19 | 0.026 |
| | MAPK signaling pathway | 3 | 46 | 0.032 |

miR=microRNA
